# Integrated multi-omics analysis identifies microbial and metabolic signatures and drivers of CNS autoimmunity

**DOI:** 10.64898/2026.01.08.698420

**Authors:** Theresa L. Montgomery, Emily A. Nelson, Lauren A. Downs, Eamonn R. Heney, Margaret Frances J. Lee, Cameron Martino, Daniel McDonald, Gibraan Rahman, Rob Knight, Dimitry N. Krementsov

**Affiliations:** Department of Biomedical and Health Sciences, University of Vermont, Burlington, VT 05401, USA; Leaven Foods Inc., San Diego, CA, 92126 USA; Department of Pediatrics, University of California, San Diego, La Jolla, CA, USA; Bioinformatics and Systems Biology Graduate Program, University of California San Diego, La Jolla, CA, USA; Department of Computer Science and Engineering, University of California San Diego, La Jolla, CA, USA; Shu Chien-Gene Lay Department of Bioengineering, University of California San Diego, La Jolla, CA, USA; Halıcıoğlu Data Science Institute, University of California, San Diego, La Jolla, CA, USA; Center for Microbiome Innovation, University of California, San Diego, La Jolla, CA, USA

**Keywords:** Multiple sclerosis, microbiome, metabolomics, multi-omics, *Limosilactobacillus. reuteri*, cresol, indole

## Abstract

Multiple sclerosis (MS) is an autoimmune disease of the central nervous system (CNS) driven by genetic and environmental determinants. The gut microbiome of people with MS (pwMS) is distinct and influences disease through immunomodulatory metabolite production. Circulating metabolites are altered in pwMS, but identifying microbial-metabolic drivers remains challenging. We previously showed that colonization by the gut commensal *Limosilactobacillus reuteri* (*L. reuteri*) exacerbates disease in the experimental autoimmune encephalomyelitis (EAE) model of MS, in a tryptophan-dependent manner. Here, we integrated microbiomic and metabolomic datasets from a longitudinal EAE study utilizing high and low tryptophan diets in mice colonized or not with *L. reuteri*. Gut microbiome dynamics under short- and long-term alterations in tryptophan bioavailability, were affected by diet, microbiome context, or disease. During short-term dietary intervention, *L. reuteri* colonization exerted a greater impact on microbiome composition than tryptophan bioavailability. With longer dietary exposure and EAE progression, high dietary tryptophan and *L. reuteri* colonization synergized to elicit profound microbiota changes, including alterations in Lachnospiraceae, *Blautia*, and *Akkermansia*. Integration of metabolomic and microbiomic datasets using joint Robust Aitchison PCA revealed clusters of associated metabolites and microbiota enriched for functional pathways, including bile acid and tryptophan metabolism. Metabolites outperformed microbiota in predicting EAE severity, identifying p-cresols and indoles as top disease-associated metabolites. Treatment with p-cresol or 3-indoleglyoxylic acid exacerbated EAE, enhanced proinflammatory T cell responses, and increased cerebellar pathology. These data demonstrate that dietary responses are shaped by gut microbiome composition and that integrated microbiomic-metabolomic analyses can identify drivers of disease worsening in MS.

**IMPORTANCE:** MS is a multifactorial disease influenced not only by genetics but also by environmental factors, potentially including diet and the composition of the gut microbiome. We show that interactions between diet and commensal gut microbiota profoundly impact levels of immunomodulatory systemic metabolites, including several that are associated with disease in pwMS. Importantly, we demonstrate that individual gut microbiota produced metabolites are sufficient to worsen disease in a mouse model of MS. Integration of gut microbiome and blood metabolite datasets combined with subsequent predictive modeling, may bolster biomarker identification and the capacity to predict disease severity in pwMS, as compared to performance of individual datasets alone. These findings highlight metabolites as key mediators linking diet and the gut microbiota to neuroinflammation. Importantly, this work suggests that targeting microbial metabolites or modifying diet–microbiome interactions may represent new strategies to reduce disease activity in MS and related autoimmune disorders.

## BACKGROUND

Multiple sclerosis (MS) is a chronic autoimmune disease of the central nervous system (CNS) and the most common cause of non-traumatic neurological disease in young adults [1, 2]. Global prevalence of MS is rising, in both developing and developed countries, affecting more than 1 million people in the United States and more than 2 million people worldwide [3, 4]. MS is an immune-mediated disease instigated by infiltration of autoreactive T cells into the CNS, initiating an inflammatory cascade leading to the recruitment of additional inflammatory immune cell subsets, persistent activation of resident microglia and macrophages, and breakdown of the blood-brain barrier [1, 2, 5]. This neuroinflammation and gliosis ultimately causes destruction of the myelin sheath and oligodendrocyte loss, leading to heterogeneous neurological symptoms and disability, ranging from visual disturbances and cognitive impairment to sensory, motor, and psychiatric symptoms [6–9].

The etiology of MS is complex and while incompletely understood, is ultimately caused by a combination of both genetic and environmental risk factors [10–14]. Genetics accounts for approximately 30% of disease risk, while the remaining 70% is due to the environment or gene by environmental interactions. Environmental risk factors for MS include reduced vitamin D intake, Epstein Barr virus (EBV) infection, low UV radiation exposure, cigarette smoking, diet, obesity, and potentially, the composition of the gut microbiome [10–12, 15]. To date, numerous studies have determined that people with MS (pwMS) have a distinct composition of the gut microbiota as compared to their healthy control counterparts [16–23]. Hallmarks of the MS specific gut microbiome include a reduction in short-chain fatty acid (SCFA)-producing microbes including *Bacteroidaceae, Ruminococcaceae* and *Clostridial clusters XIVa* and *IV* [21, 24, 25], altered levels of *Akkermansia muciniphila* (*A. muciniphila*), diminished levels of *Prevotella* species, and alterations in *Parabacteroides*, *Blautia*, and *Dorea*, as well as increases in *Actinobacteria* and genera within that phylum including *Bifidobacterium*, *Streptococcus*, *Desulfovibrionaceae*, and *Coriobacterium* [16–19, 24–31].

The gut microbiome is postulated to impact CNS autoimmunity through production of immunomodulatory metabolites that affect immune cell function locally in the gut, or enter systemic circulation to impact distal sites, including the CNS [32]. Consistent with observed alteration of the MS gut microbiome, fecal, serum, and plasma metabolites are also altered in pwMS including a reduction in bile acids [33, 34] and short-chain fatty acids (SCFAs) [35–42], as well as altered levels of tryptophan associated metabolites [43–54] and cresols [48, 54].

A handful of studies to date have leveraged a multi-omics approach in pwMS to determine how gut microbiome compositional changes correlate with immunological and metabolomic differences. In a six-month longitudinal study looking at microbiome, immune profiling, and serum metabolites, methionine metabolism was linked to an increased Th17 response and *Bacteroides thetaiotaomicron* [39]. Fitzgerald et al. paired serum or plasma global metabolomics with immunological profiling, identifying an alteration in aromatic amino acid metabolism defined by an increase in the metabotoxins, p-cresol glucuronide and p-cresol sulfate, and an decrease in lactate metabolites (indole lactate, and imidazole lactate), which were associated with altered expression of genes related to aromatic amino acid metabolism in circulating monocytes [48]. Interestingly, a separate study also identified reduced levels of indole lactate as well as indole propionate in circulation, correlating with a reduction in the microbiota that produce them as well as a reduction in SCFA-producing microbiota [49]. In relapsing-remitting MS (RRMS), increases in the *Slackia* and *Lactobacillus* genus have been linked to altered levels of glycerophospholipid associated metabolites [55] and a reduction in SCFA-producing microbiota was linked to reduced levels of fecal butyrate and propionate [37]. The international MS Microbiome Study (iMSMS) profiled fecal and serum metabolomics in addition to the composition of the gut microbiome [16]. Interestingly, while noting a decrease in prominent SCFA-producing gut microbiota including *Faecalibacterium prausnitzii* and *Blautia* species, lower levels of fecal SCFAs predominantly were associated with disease modifying therapies but not disease status per se. These studies highlight the utility of a multi-omics approach to define the functional implications of the gut microbiota in mediating disease. However, experimental studies are needed to functionally demonstrate the link between specific microbes, metabolites, and disease outcomes.

We and others have previously shown that colonization by the gut commensal bacterium *L. reuteri* is sufficient to exacerbate CNS autoimmune disease in the primary autoimmune model of MS, experimental autoimmune encephalomyelitis (EAE) [56–58]. In subsequent studies, we demonstrated that this response is dependent on the availability of host dietary tryptophan [56, 59]. Using a combination of *in vitro* and *in vivo* metabolomics, we have shown that *L. reuteri* produces a wide array of tryptophan-dependent metabolites, including cresols and indoles *in vitro*, while *in vivo* colonization alters the systemic metabolic profile in a tryptophan-dependent manner. Furthermore, we have shown that *L. reuteri* colonization under tryptophan-sufficiency remodels the global composition of the gut microbiome, leading to depletion in SCFA-producing gut microbiota [60]. Mechanistically, these data suggest that *L. reuteri* drives CNS autoimmunity via a combinatorial effect on the gut microbiome composition and alteration of systemic metabolites. However, the interaction between these two divergent mechanisms and the functional requirements for EAE exacerbation remain unclear.

In a longitudinal study, we examined the gut microbiome and systemic metabolites in mice exposed to high- or low-tryptophan diets, with or without *L. reuteri* colonization. This approach revealed key baseline microbiota characteristics influenced by diet, microbiome context, and disease progression. In the short-term dietary intervention, microbiota composition was more strongly influenced by *L. reuteri* colonization than by tryptophan bioavailability. During extended dietary intervention and EAE progression, high tryptophan intake and *L. reuteri* colonization interacted to drive substantial microbiota shifts, including alterations in *Lachnospiraceae*, *Blautia coccoides*, and *Akkermansia muciniphila*. Integrating serum metabolomics with microbiome data revealed distinct clusters of closely linked metabolites and microbes, enriched in metabolic pathways that included bile acid, nucleotide, nicotinate/nicotinamide, and tryptophan metabolism. Using Random Forest modeling, we found that metabolites were stronger predictors of EAE severity than was the microbiota, highlighting p-cresols and indoles as key biomarkers of disease. Administration of p-cresol or 3-indoleglyoxylic acid, acid in the EAE model worsened disease severity, promoting proinflammatory T cell responses in the CNS, and increased demyelination. These data highlight the utility of multi-omics approaches to identifying the microbial and metabolic drivers of disease worsening in MS and suggest that dietary response is modulated by the composition of the baseline gut microbiome.

## RESULTS

### L. reuteri colonization impacts microbial dynamics more than short-term dietary tryptophan intervention

In our previous studies, we found that stable colonization by *L. reuteri* exacerbates EAE [56]. Moreover, we have shown that this effect is dependent on the availability of host dietary tryptophan [59]. Here, we set out to determine how tryptophan bioavailability impacts the microbiome, metabolome, and severity of EAE, and whether experimental colonization by specific commensal microbial species modulated these effects. First, to determine the impact of dietary tryptophan bioavailability on gut microbiome composition, we leveraged our previously established gut microbiota transplantation and vertical transmission model [59]. Specifically, we previously colonized germ-free (GF) C57BL/6 (B6) mice with cryopreserved stocks of 1) normal B6 cecal microbiota (naturally devoid of *L. reuteri*) or 2) normal B6 cecal microbiota supplemented with 10^9^ CFU *L. reuteri*, to establish G_0_ (generation zero) breeding pairs. G_0_ breeding pairs vertically passed microbiota to their offspring (G_1_). At 7 weeks of age, G_1_ offspring harboring B6 or B6 + *L. reuteri* microbiota were randomized to either a 0.02% (low) or 0.8% (high) tryptophan (Trp) diet 1 week prior to induction of EAE, and maintained on these diets for a full 30-day EAE disease course (**Fig. 1**). We collected fecal samples for full-length 16S (FL16S) sequencing to interrogate the gut microbiome composition at two distinct timepoints: 1) 1-week post dietary intervention (Week 1, prior to disease induction) and 2) 5-weeks post dietary intervention (Week 5, following a full 30-day disease course) (**Fig. 1**).

**Figure 1.**
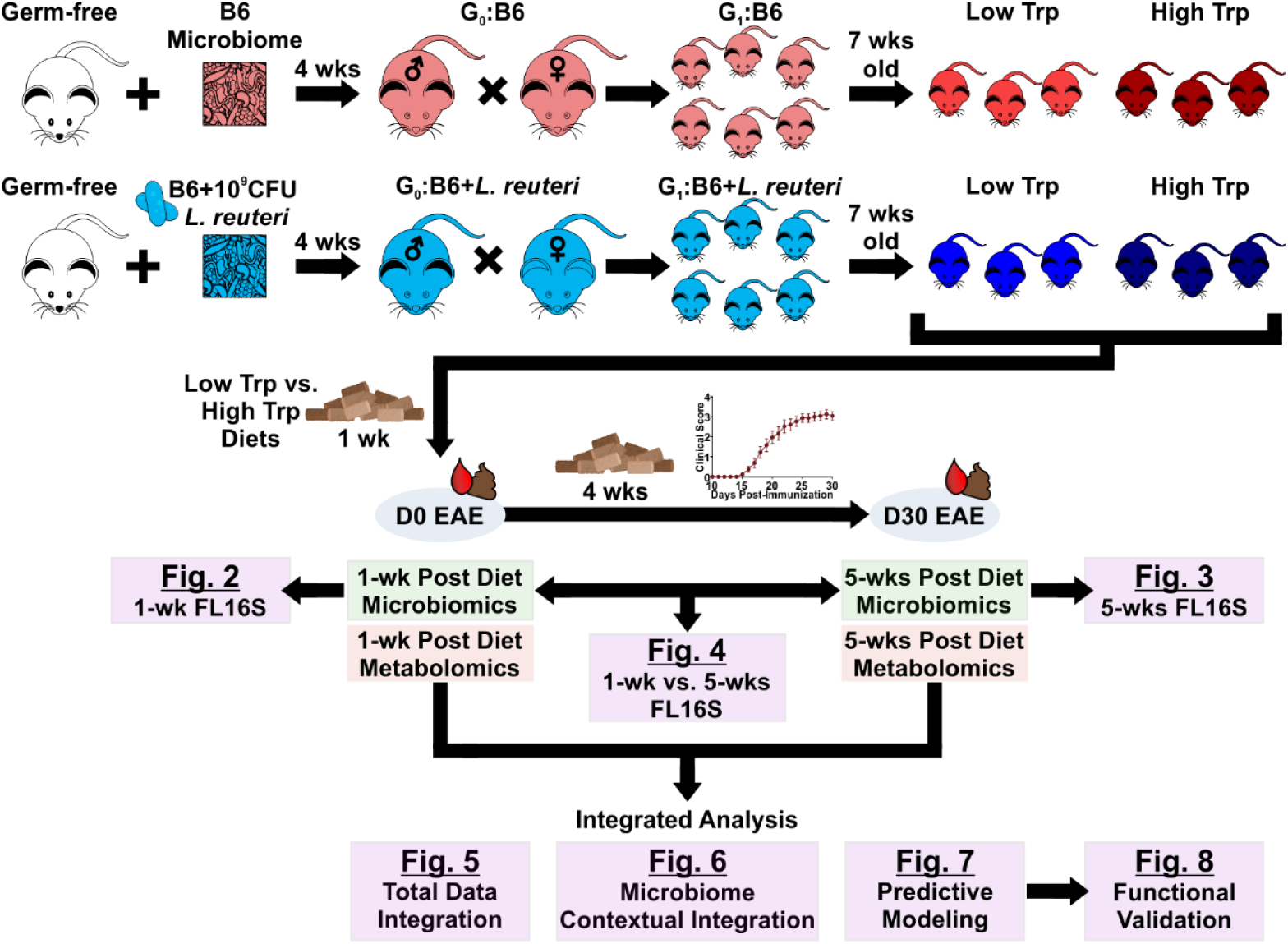
Schematic of microbiome transplantation, vertical transmission, and dietary tryptophan modulation model, depicting timeline for fecal and serum collection. Cryopreserved B6 cecal gut microbiome, naturally lacking *L. reuteri* (B6) or supplemented with 10^9^ CFU *L. reuteri* (B6 + *L. reuteri*) was introduced as a one-time inoculation into germ free B6 breeding pairs, which were used to generate experimental offspring that were randomized to high or low tryptophan (Trp) diets. Fecal samples were collected at 1 week post dietary intervention and interrogated by full-length 16S DNA sequencing (FL16S), followed by compositional analyses (see Methods).

To elucidate the proximal effect of dietary tryptophan on gut microbiome composition, we initially limited our analysis to the week 1 timepoint. To determine the broad effect of dietary tryptophan on each gut microbiome configuration, we first analyzed metrics of alpha and beta diversity, binning experimental groups by diet and *L. reuteri* status as follows: B6 Low Trp, B6 + *L. reuteri* Low Trp, B6 High Trp and B6 + *L. reuteri* High Trp. Alpha diversity, as measured by Shannon’s diversity index, a metric of intra-individual gut microbial diversity, was similar when compared between *L. reuteri* contexts (B6 vs. B6 + *L. reuteri*) and between diets (low or high tryptophan) (**Fig. 2A**). We have previously reported that colonization of the gut by *L. reuteri* remodels the overall composition of the gut microbiome [60]. Consistently, beta diversity, as measured by unweighted Unifrac distance, a metric of inter-individual gut microbial diversity, was significantly different by microbiome (p=0.001; PERMANOVA) (**Fig. 2B** and **C**). Interestingly, neither diet (tryptophan bioavailability) nor the interaction between diet and microbiome, significantly influenced beta diversity at this timepoint (**Fig. 2B** and **C**). Exploratory analysis of the top 11 most abundant genera, suggested that consistent with beta diversity analyses, *L. reuteri* colonization significantly remodeled the gut microbiome in the context of both high and low dietary tryptophan, leading to increased *Akkermansia* and *Turicibacter* at the expense of the *Faecalibaculum* genus (**Fig. 2D**). To statistically evaluate specific compositional differences by gut microbiome and diet, we next analyzed changes at each taxonomic rank using linear models for differential abundance analysis (LinDA) [61], comparing either 1) the B6 vs. B6 + *L. reuteri* gut microbiomes, adjusted for effect of diet, or 2) Low vs. high tryptophan diets, adjusted for effect of microbiome group. Analysis with LinDA revealed that experimental microbiome exerted a strong effect at this timepoint, with the presence of *L. reuteri* resulting in increased *Akkermansia muciniphila* (*A. muciniphila*)*, Lactobacillus johnsonii* (*L. johnsonii*), and amplicon sequence variants (ASVs) of the *Turicibacter* and *Roseburia* genus and *Muribaculaceae* family, as well as modulation of *Lachnospiraceae* and *Oscillibacter* members (**Fig. 2E**). Notably, diet, when adjusted for microbiome configuration, did not lead to significant alteration of any individual ASVs (**Fig. 2C**), consistent with the PERMANOVA results.

**Figure 2.**
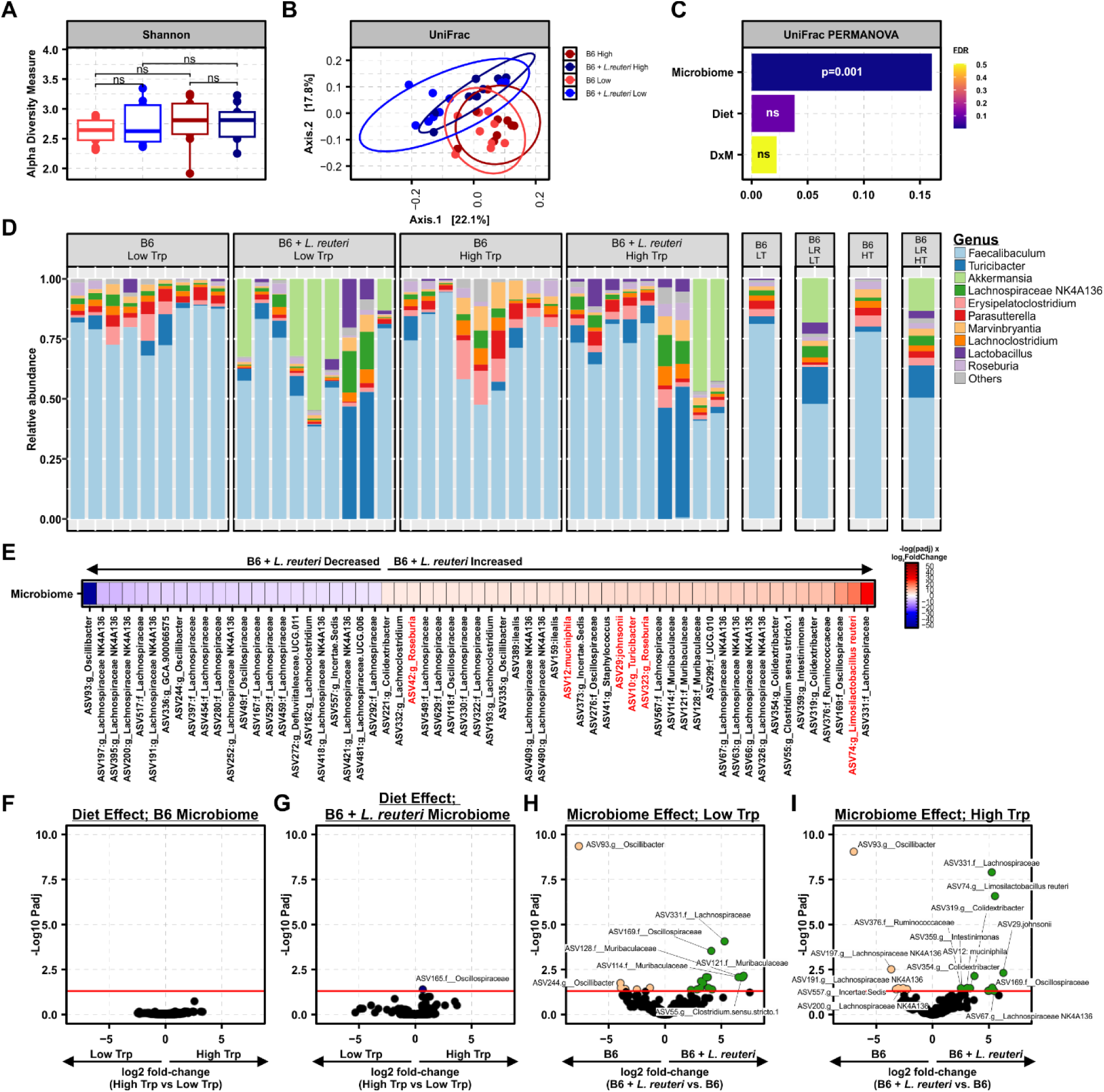
*L. reuteri* colonization exerts a stronger influence on microbial composition than short-term dietary tryptophan modulation. Mice colonized by B6 or B6 + *L. reuteri* microbiomes were randomized to high or low Trp diets, followed by EAE induction (see Fig. 1). Fecal samples were collected at 1 week post dietary intervention (directly prior to EAE induction) and interrogated by full-length 16S DNA sequencing, followed by compositional analyses. (**A**) Alpha (Shannon), (**B**) beta (unweighted UniFrac) and (**C**) unweighted UniFrac PERMANOVA diversity analysis in each gut microbiome × diet treatment group, with significance determined using Wilcoxon rank sum non-parametric (**A**) or Adonis (**B**) testing. (**D**) Stacked bar plots of top 11 most abundant genera as a proportion of total reads. (**E**) Differentially abundant ASVs (represented as a taxonomic best-hit) between the B6 and B6 + *L. reuteri* gut microbiomes as determined by LinDA when adjusting for diet, using a cutoff of p_adj_ ≤ 0.05. Warm colors represent increased abundance in the presence of *L. reuteri* and cooler colors represent decreased abundance, scaled by the -log(p_adj_) × log2 fold-change. Differentially abundant ASVs driven by diet in mice colonized by the B6 (**F**) or B6 + *L. reuteri* (**G**) gut microbiomes, where log2 fold-change indicates increased abundance with a high tryptophan diet when positive and decreased abundance when negative, as determined with LinDA, using a cutoff of p_adj_ ≤ 0.05. Differentially abundant ASVs in low tryptophan (**H**) or high tryptophan (**I**) fed mice driven by microbiome where log fold-change indicates increased abundance within the B6 + *L. reuteri* gut microbiome when positive and decreased abundance when negative as determined using LinDA with a cutoff of padj ≤ 0.05.

To determine the impact of diet in each gut microbiome context, we evaluated differential ASV abundance between high and low tryptophan diets subset to either the B6 or B6 + *L. reuteri* gut microbiome (**Fig. 2F** and **G**). Consistent with alpha and beta diversity analyses, there were no significant ASV level differences as a function of diet in the B6 gut microbiome, and only a single ASV that was modestly increased by high tryptophan diet in the B6 + *L. reuteri* microbiome (**Fig. 2F** and **G**). Next, to evaluate the effect of microbiome in each dietary context, we compared the effect of *L. reuteri* status on ASV level abundance, subset to either low or high tryptophan diet (**Fig. 2H** and **I**). Consistent with combined analysis adjusted for diet and microbiome (**Fig. 2E**), numerous ASVs were significantly altered with *L. reuteri* status. While *L. reuteri* colonization in the context of low dietary tryptophan led to alteration of *Oscillospiraceae* and an increase in *Muribaculaceae* (**Fig. 2H**), *L. reuteri* colonization in the context of a high tryptophan diet resulted in an increase in *L. johnsonii* and ASVs in the *Colidextribacter* genus, with a notable decrease in several ASVs belonging to the *Lachnospiraceae NK4A136* genus (**Fig. 2I**), known producers of SCFAs [60]. Taken together, these data suggest that 1 week following post-dietary intervention, tryptophan bioavailability had a minimal effect on the gut microbiome, while the presence of *L. reuteri* was sufficient to alter gut microbiome composition, in a manner that differed depending on the presence of tryptophan.

### Modification of dietary tryptophan bioavailability during EAE progression alters gut microbiome composition in a microbiota composition-dependent manner

To assess the interplay between diet and gut microbiome context following EAE initiation and a longer dietary exposure, we next evaluated the composition of the gut microbiome at the 5-week post-diet timepoint following a full 30-day disease course (**Fig. 1**). Shannon’s diversity index was significantly different as a function of diet in both B6 and B6 + *L. reuteri* microbiomes (p=0.02 and p=0.03 respectively), and as a function of microbiome in the context of a high tryptophan diet (p=0.03) (**Fig. 3A**). Beta diversity, as assessed by unweighted Unifrac distance, was also significantly different as a function of diet, microbiome, and diet by microbiome interaction (p=0.001, p=0.001, and p=0.007, respectively) (**Fig. 3B** and **C**). These data suggest that, by the 5-week timepoint, diet had a more profound impact on gut microbiome composition than observed at the 1-week timepoint, and further that diet has a more profound impact on gut microbiome composition in the context of *L. reuteri* colonization. Consistent with diversity analyses, visualization of the top 11 genera suggested that both diet and the presence of *L. reuteri* was sufficient to alter the composition of the gut microbiome. While a high tryptophan diet led to an expansion of *Parasutterella* in the B6 gut microbiome, an expansion of *Blautia* and *Roseburia* were observed in the B6 + *L. reuteri* microbiome (**Fig. 3D**).

**Figure 3.**
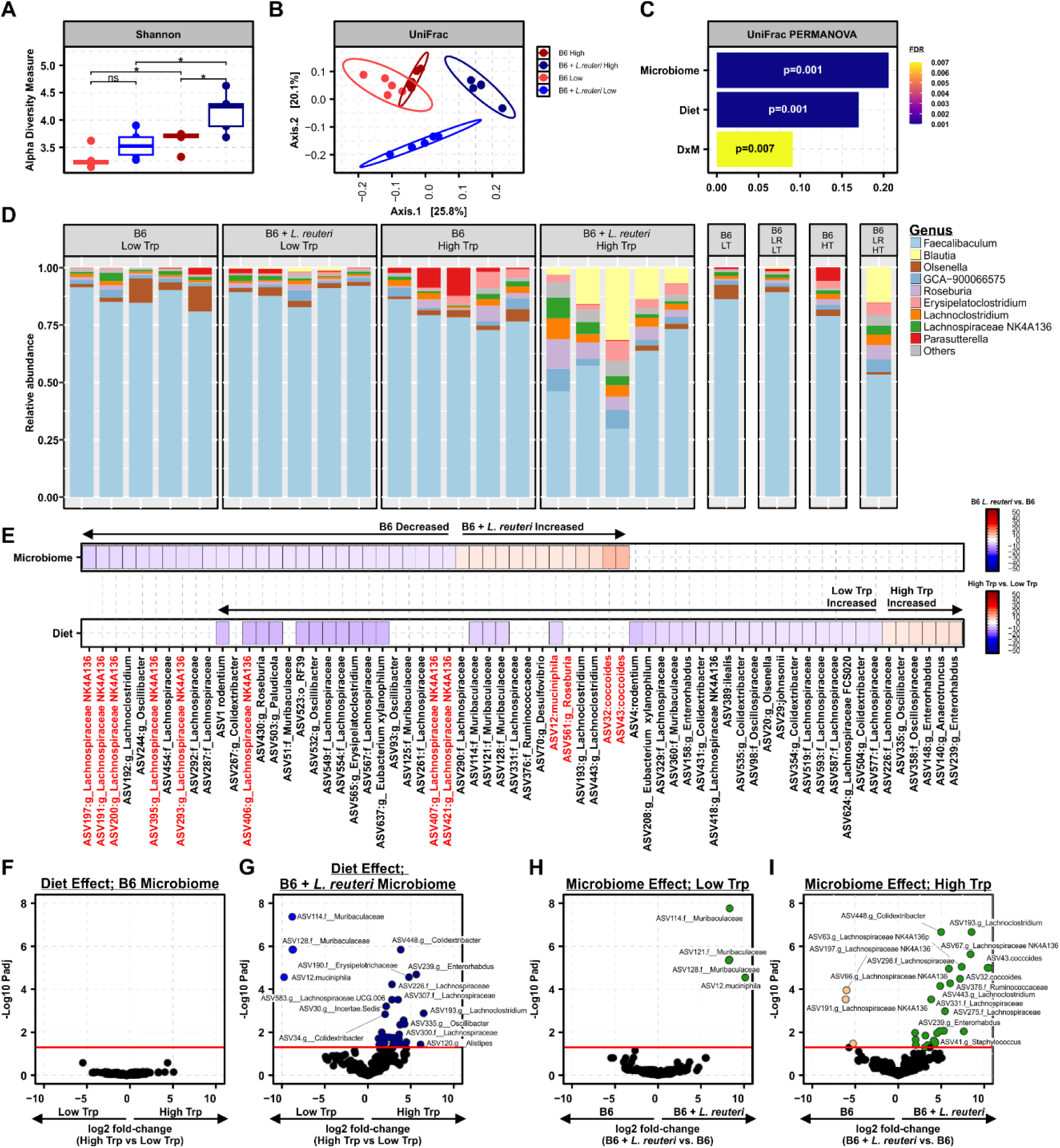
Baseline microbiota influence the gut microbial compositional response to prolonged changes in tryptophan bioavailability. Mice colonized by B6 or B6 + *L. reuteri* microbiomes were randomized to high or low Trp diets, followed by EAE induction (see Fig. 1). Fecal samples were collected at 5 weeks post dietary intervention (30 days post EAE induction) and interrogated by full-length 16S DNA sequencing, followed by compositional analyses. (**A**) Alpha (Shannon), (**B**) beta (unweighted UniFrac) and (**C**) unweighted UniFrac PERMANOVA diversity analyses in each gut microbiome × diet treatment group, with significance determined using Wilcoxon rank sum non-parametric (**A**) or Adonis testing (**B**). (**D**) Stacked bar plots of top 11 most abundant genera as a proportion of total reads. (**E**) Differentially abundant ASVs (represented as a taxonomic best-hit) between the B6 and B6 + *L. reuteri* gut microbiomes when adjusted for diet (top), or between low and high tryptophan fed mice when adjusted for gut microbiome (bottom), as determined by LinDA, using a cutoff of p_adj_ ≤ 0.05. Warm colors represent increased abundance and cooler colors represent decreased abundance, scaled by -log10(padj) × log2 fold-change. Differentially abundant ASVs driven by diet in the B6 (**F**) or B6 + *L. reuteri* (**G**) gut microbiomes, where log fold-change indicates increased abundance with a high tryptophan diet when positive and decreased abundance when negative, as determined using LinDA, with a cutoff of p_adj_ ≤ 0.05. Differentially abundant ASVs driven by microbiome in low tryptophan (**H**) or high tryptophan (**I**) fed mice, where log fold-change indicates increased abundance within the B6 + *L. reuteri* gut microbiome when positive and decreased abundance when negative as determined using LinDA with a cutoff of padj ≤ 0.05.

To isolate the gut microbiome as an experimental variable, differential abundance of individual ASVs was analyzed using LinDA, adjusting for diet. Colonization with *L. reuteri* led to an expansion of *A. muciniphila*, two *Blautia coccoides* ASVs, and a marked depletion of ASVs belonging to the SCFA-producing *Lachnospiraceae NK4136* genus (**Fig. 3E**). Similarly, to evaluate the impact of diet, independent from microbiome context, differential abundance of individual ASVs was analyzed using LinDA, adjusting for microbiome. There was low overlap between the effect of microbiome and effect of diet, with a total of 11 ASVs displaying uniform directionality (depletion as a function of high dietary tryptophan and the presence of *L. reuteri*). Interestingly, *A. muciniphila* abundance was reduced by provision of a high tryptophan diet, as were several *Lachnospiraceae* and *Colidextribacter* members (**Fig. 3E**).

To assess the impact of diet within each gut microbiome context, we next subset our analysis of the effect of diet to the B6 or B6 + *L. reuteri* gut microbiome. There were no significant changes of individual ASVs caused by diet in mice colonized by the B6 gut microbiome (**Fig. 3F**). In contrast, a high tryptophan diet led to a depletion of *A. muciniphila* and *Muribaculaceae* members, with an expansion of *Alistipes*, *Lachnospiraceae*, and *Colidextribacter* members in the context of the B6 + *L. reuteri* gut microbiome (**Fig. 3G**). To evaluate the impact of each experimental gut microbiome within each diet, we next subset our analysis to include either low or high tryptophan diet fed mice. Consistent with diversity analyses, there were minimal changes caused by *L. reuteri* colonization in the context of a tryptophan depleted diet (**Fig. 3H**). Meanwhile, *L. reuteri* colonization in the context of a tryptophan-rich diet led to a significant depletion of SCFA-producing *Lachnospiraceae NK4136* members, with an expansion of *Blautia coccoides*, *Lachnoclostridium,* and several other ASVs (**Fig. 3I**). These data suggest that EAE progression combined with prolonged dietary exposure has a more substantial impact on gut microbiome composition, but only in the presence of *L. reuteri* colonization. Conversely, high dietary tryptophan promoted a much more profound effect of *L. reuteri* colonization on the overall gut microbiota composition. Both findings are consistent with the well-known capacity of *L. reuteri* to catabolize tryptophan [59], and suggest that the effect of diet is highly dependent on the baseline composition of the gut microbiota.

### Tryptophan bioavailability and L. reuteri colonization alter gut microbiome temporal dynamics

To characterize the specific changes in the gut microbiota over time, during extended exposure to a diet replete with or devoid of tryptophan and throughout the course of EAE, we next compared a subset of matched fecal samples collected from the same mice at week 1 and week 5 post-dietary intervention (**Fig. 1**). The B6 gut microbiome in the context of a low tryptophan diet displayed a reduction in alpha diversity over time as measured by Shannon’s index (p=0.04), which was not seen in the context of the high tryptophan diet (**Fig. 4A**). Meanwhile, the B6 + *L. reuteri* gut microbiome in the context of a high tryptophan diet displayed significantly increased alpha diversity over time (p=0.004), that was again absent in the context of low tryptophan (**Fig. 4A**). To determine if the variation in alpha diversity over time significantly differed between diet and gut microbiome contexts, change in alpha diversity over time was represented as fold-change, and the magnitude of this change was compared between specific groups of interest. A significant difference in the change in alpha diversity over time was observed comparing high and low tryptophan fed mice harboring both microbiomes (p=0.03 and p<0.001, B6 and B6 + *L. reuteri*, respectively) (**Fig. 4A**). Comparing between the two gut microbiomes in the context of high dietary tryptophan also demonstrated a significant difference in the change in alpha diversity over time (p=0.04), not observed in the context of a low tryptophan diet (p=0.64) (**Fig. 4A**). These data suggest that high tryptophan promotes more dynamic changes in gut microbial richness than does low tryptophan. Analysis of beta diversity (unweighted Unifrac) revealed that no significant compositional differences over time occurred in the B6 microbiome regardless of diet, while in both dietary contexts (low and high tryptophan) the B6 + *L. reuteri* microbiome composition changed over time, with a more pronounced change with provision of a tryptophan replete diet (R^2^=0.27 vs. R^2^=0.50) (**Fig. 4B**). These data suggest that the colonization by *L. reuteri* promotes more dynamic changes in gut microbiome composition over the course of extended dietary exposure and EAE progression, particularly in the context of high tryptophan.

**Figure 4.**
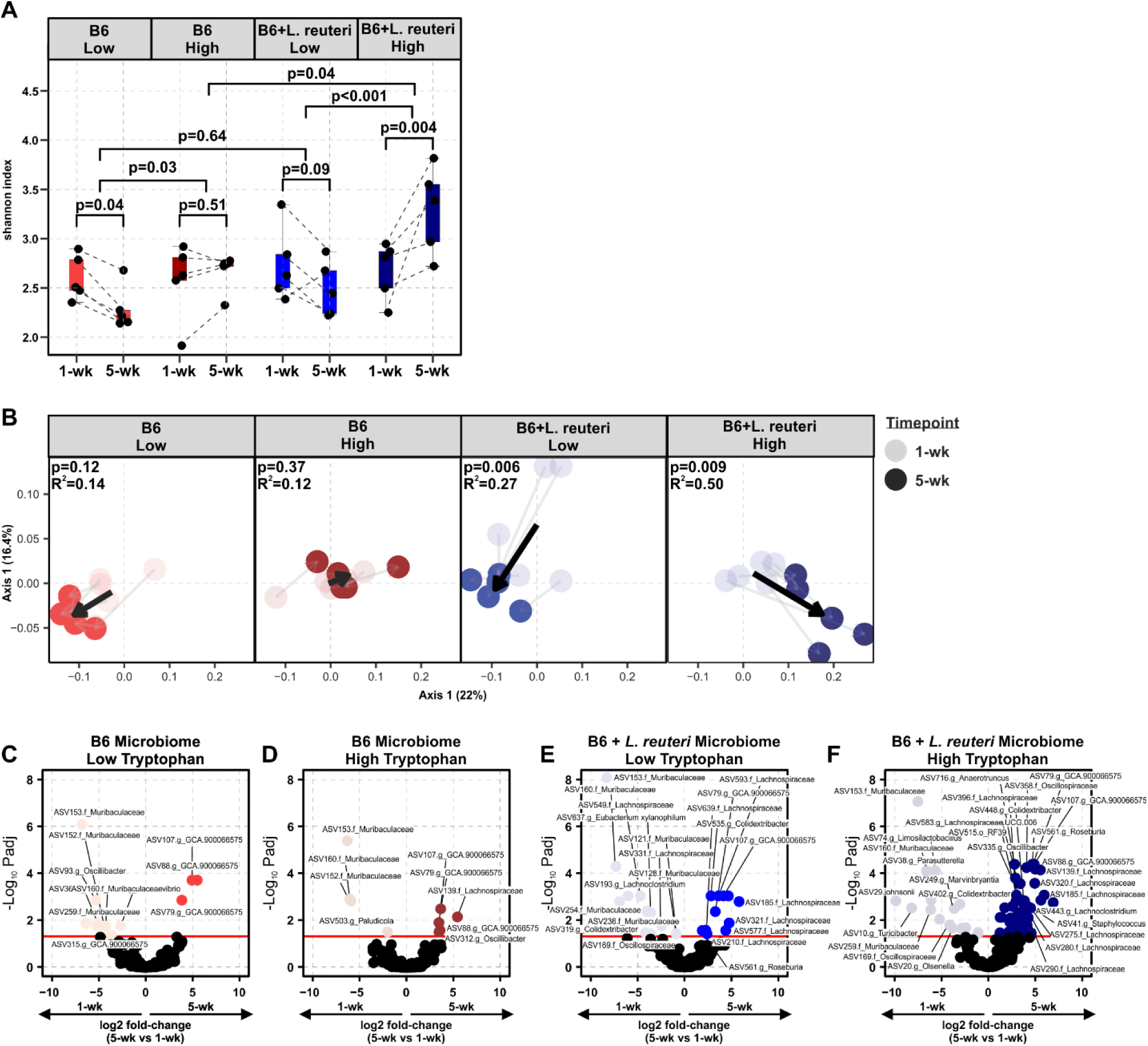
Gut microbiome temporal dynamics are shaped by tryptophan bioavailability and *L. reuteri* colonization. Paired microbiome samples collected from mice colonized by B6 or B6 + *L. reuteri* microbiomes on high and low Trp diets at 1 week and 5 weeks post-dietary intervention were analyzed for the effect of time (see Fig. 1). (**A**) Alpha (Shannon) and (**B**) beta (unweighted UniFrac) diversity analysis in each gut microbiome × diet treatment group at the 1-wk and 5-wk post-dietary intervention timepoints. Alpha diversity between timepoints within group is a paired analysis of the absolute change between timepoints using linear models and a t-test for significance. Difference in the magnitude of change in alpha diversity over time between experimental diet-by-microbiome groups represents log-fold change between groups with significance determined by t-test. Beta diversity statistical significance of change over time in each group was evaluated with adonis testing. Differentially abundant ASVs by time in the B6 gut microbiome in mice fed a low (**C**) or high (**D**) tryptophan diet, where log fold-change indicates increased abundance at the 5-wk timepoint when positive and decreased abundance when negative, as determined using LinDA with a cutoff of p_adj_ ≤ 0.05. Differentially abundant ASVs by time in the B6 + *L. reuteri* gut microbiome in mice fed a low (**E**) or high (**F**) tryptophan diet, where log fold-change indicates increased abundance at the 5-wk timepoint when positive and decreased abundance when negative, as determined using LinDA using a cutoff of p_adj_ ≤ 0.05.

To assess the dynamics of individual species over time as a function of diet and gut microbiome context, we analyzed ASV differential abundance between 1 and 5 week time points using LinDA, in each experimental group. Notably, mice harboring in the B6 gut microbiome, fed either a low or high tryptophan diet, exhibited a limited number of changes by time, consisting of a depletion in *Muribaculaceae* and increased abundance of the genus *GCA_900066575* (*Lachnospiraceae*) (**Fig. 4C** and **D**). In contrast, mice colonized with the B6 + *L. reuteri* gut microbiome displayed a greater number of ASV-level changes over time, especially when fed a high tryptophan diet (B6: low Trp =12, high Trp =9, vs. B6 + *L. reuteri*: low Trp = 24, high Trp =71; **Fig. 4E** and **F**). In the context of depletion of tryptophan and the B6 + *L. reuteri* microbiome, time resulted in an alteration of *Lachnospiraceae*, *Muribaculaceae*, and *Colidextribacter* species, as well as an increase in *Roseburia* and decrease in *Eubacterium xylanophilum* (**Fig. 4E**). Meanwhile, in mice colonized by the B6 + *L. reuteri* microbiome and fed a tryptophan replete diet, time led to increased *Colidextribacter*, *Blautia coccoides, Enterococcus faecalis*, *Staphylococcus xylosus*, *Tuzzerella, Roseburia*, several *Ruminococcaceae, Oscillibacter*, and *Lachnospiraceae* members, with a decrease *L. johnsonii*, *Faecalibaculum rodentium*, *Turicibacter*, and *Parasutterella* ASVs, and, surprisingly, a decrease in *L. reuteri* itself (**Fig. 4F**). Together with the beta diversity analyses, these data suggest that the *L. reuteri*-colonized gut microbiome is more dynamic during extended dietary exposure and the course of EAE than is a microbiome lacking this species. Moreover, these dynamics are further promoted by the bioavailability of dietary tryptophan.

### Multiomic integration reveals functionally distinct microbe:metabolite associations

We had previously characterized the global impact of tryptophan bioavailability on host circulating metabolic profiles via UPLC-MS/MS, analyzing serum collected from B6- or B6 + *L. reuteri-*colonized mice fed either the low or high tryptophan diet after 1 week and 5 weeks (following a full 30-day EAE course) (**Fig. 1**) [59]. To identify the microbiota associated with and possibly driving these observed metabolic changes, we integrated these metabolomic data with our FL16S sequencing dataset using joint robust Aitchison principal component analysis (jRPCA) [62, 63]. jRPCA is specifically tailored to the integration of multiple omics layers, leveraging an unsupervised clustering method for sparse compositional datasets such as microbiomes. We initially analyzed our total dataset consisting of matched fecal FL16S taxonomic data and serum metabolomics from all timepoints, microbiomes, and diets. jRPCA generates a feature table containing the contribution of each feature to principal components that segregate samples, which can be projected as a bi-plot. Ordination of integrated datasets along the first two principal components revealed three prominent microbe:metabolite clusters, consisting of bile acids, purine nucleotides, or indoles/cresols (tryptophan derived metabolites) and specific microbiota ASVs that were most closely associated with them, such as ASVs mapping to *GCA-900066575* (*Lachnospiraceae*), *A. muciniphila,* and *Anaerotruncus* (*Oscillospiraceae*), respectively (**Fig. 5A**). We next analyzed microbe:metabolite pairwise correlations, as identified by jRPCA. Hierarchical clustering of microbial:metabolite pairwise correlations by Euclidean distance and complete linkage followed by partitioning of the resulting row and column dendrograms, defined a total of seven distinct clusters of metabolites and eleven microbial clusters (**Fig. 5B**). Extraction of the metabolites within each cluster and pathway enrichment analysis revealed enrichment of distinct metabolic pathways for each cluster, including significant enrichment of alpha linolenic acid and linoleic acid metabolism in cluster 1, phosphatidylcholine biosynthesis in cluster 2, amino acid biosynthesis and bile acid biosynthesis in cluster 3, glycine and serine metabolism, ammonia recycling, and purine metabolism in cluster 4, glycerophospholipid metabolism in cluster 5, pyrimidine metabolism in cluster 6, and both tryptophan metabolism and nicotinate/nicotinamide metabolism in cluster 7 (**Fig. 5C**). To explore ASV:metabolite interactions, highly correlated features (>|0.99| correlation coefficient) within each cluster were plotted as networks, where nodes represented individual species or metabolites, and edge width indicated strength of association, with positive interactions in blue and negative in red (**Fig. 5D** and **E**). Using this approach, we identified 16 ASVs to be most strongly correlated with bile acid metabolism, mainly belonging to the *Clostridia* class including *Lachnospiraceae*, *Lachnospiraceae NK4A136*, *Oscillospiraceae,* as well as *Olsenella* and *Muribaculaceae* members. Interestingly, ASV74, representing *L. reuteri*, was placed in a sub-cluster of prominent bile acids with high connectivity (**Fig. 5D**). Within cluster 7, which was enriched for tryptophan and nicotinate & nicotinamide metabolism, a greater diversity of microbiota was represented, including *Bacteroidia* (predominately *Muribaculaceae*) and *Bacilli*, most notably, *L. reuteri*, *L. johnsonii*, *Erysipelatoclostridium*, and *Turicibacter* species, as well as various *Clostridia* species (**Fig. 5E** and **F**). Taken together, these analyses integrate microbial and metabolite features in this complex dataset to identify major sources and patterns of variance, as well as novel relationships among microbes and metabolites.

**Figure 5.**
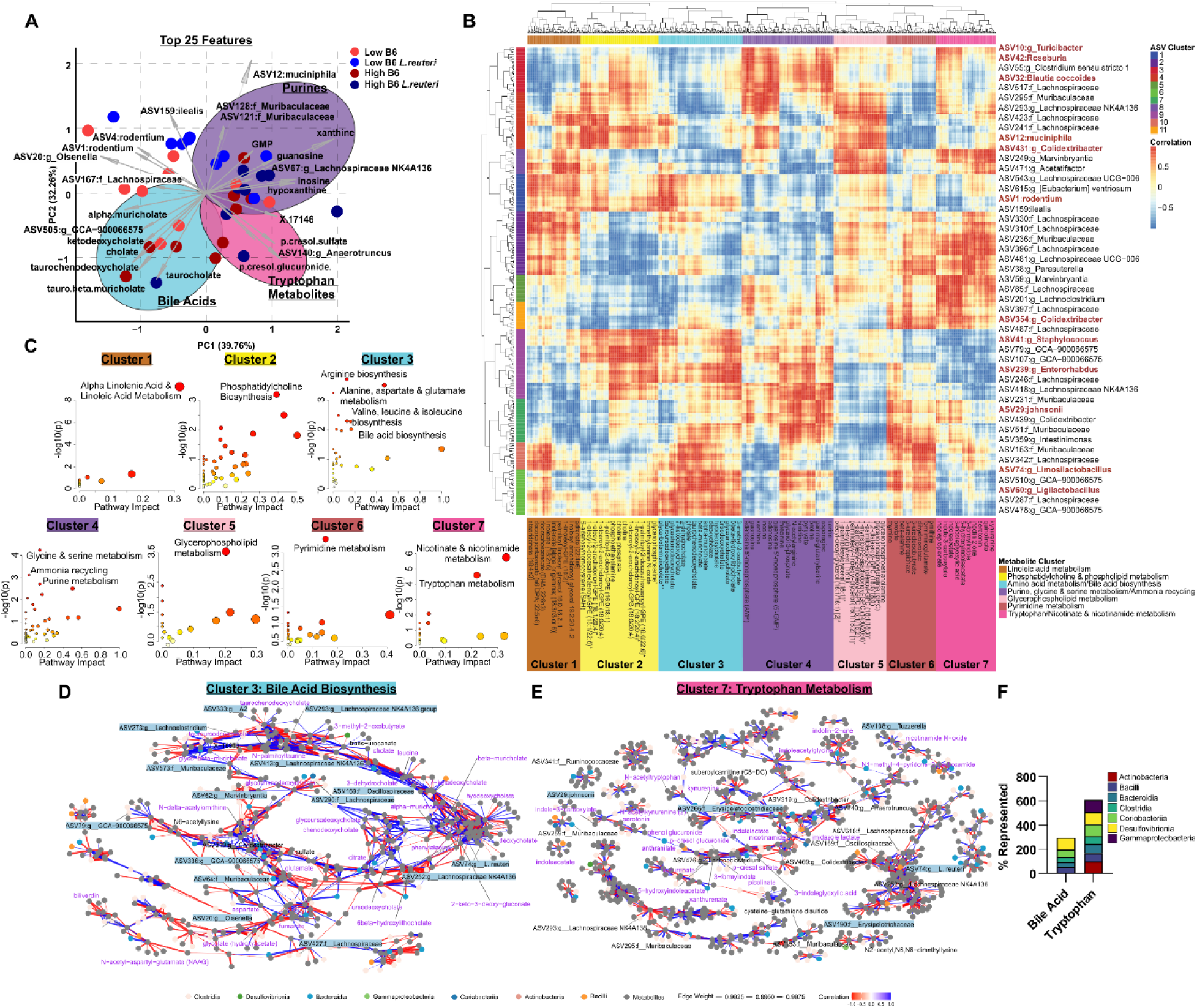
Integrated microbiomic and metabolomic analysis reveals functionally distinct clusters of microbes and metabolites. Datasets containing matched fecal microbiome and blood metabolome samples collected at 1 wk and 5 wks post-dietary intervention were integrated using joint robust principal component analysis (jRPCA), which was used to determine sample ordination in principal coordinate space and feature-feature correlation coefficients (see Methods). (**A**) Emperor biplot of integrated sample clustering in principal coordinate space based on multiomic variation, with arrows indicating the vectors of the top 25 features (microbe ASVs or metabolites) contributing to principal components. (**B**) Heatmap of correlation coefficients between ASVs (rows) and metabolites (columns), clustered using Euclidean distance and complete linkage, with clusters identified by cutting the trees at fixed heights, resulting in 7 distinct metabolite clusters and 11 ASV clusters. (**C**) Metabolites in each cluster in (**B**) were analyzed and annotated by pathway enrichment analysis, with significant enrichment at cutoff of p_adj_ ≤ 0.05. Networks of correlated ASVs and metabolites within the bile acid (**D**) or tryptophan (**E**) clusters. Data was filtered at a threshold of |0.99| strength of correlation, nodes represent metabolites (grey) or ASVs (with color assignment based on taxonomic class), edge widths represent the strength of correlation, and colors indicate the direction of correlation. (**F**) ASVs represented in the bile acid or tryptophan networks within each taxonomic class as a percentage of all ASVs in a given class in the total dataset.

### Bile acids, tryptophan metabolites, and associated microbiota robustly delineate groups by dietary context and time

To define differences in the microbial and metabolic signatures influenced by diet or time, and specific to each gut microbiome context, we next subset our analyses by gut microbiome configuration (B6 and B6 + *L. reuteri*) followed by integration of omics datasets using jRPCA. Samples segregated broadly by dietary context and timepoint along the first two principal components in both the B6 (**Fig. 6A**) and B6 + *L. reuteri* (**Fig. 6B**) gut microbiomes. Extracting the top 50 features contributing most significantly to separation along each principal component identified differences in the microbial and metabolic signatures driven by diet and time between microbiomes. In the context of the B6 gut microbiome, high tryptophan was associated with a purine nucleotide signature (guanosine, xanthine, hypoxanthine, GMP, inosine, and adenosine), indole acrylate, and several *Lachnospiraceae*, while low tryptophan was associated with bile acids (taurochenodeoxycholate, tauro-beta-muricholate, taurocholate, tauroursodeoxycholate), as well as *Roseburia* and *Blautia* genera, and interestingly *Ligilactobacillus murinus* (**Fig. 6C**), a *Lactobacillaceae* family member that shares metabolic features with *L. reuteri*, yet behaves very distinctly in terms of modulation of gut microbiota composition and EAE severity [60]. By contrast, features most significantly contributing to separation of samples by diet in the B6 + *L. reuteri* microbiome included metabolites associated with tryptophan metabolism, where high dietary tryptophan was associated with kynurenate, quinolinate, and xanthurenate, as well as the p-cresols: p-cresol-glucuronide and p-cresol-sulfate (**Fig. 6E**), putative microbial tyrosine metabolites that we previously determined to be tryptophan dependent [59]. Similar to the B6 microbiome, low dietary tryptophan was associated with an increase in bile acids (hypotaurine and alpha-muricholate), as well as uniquely associated with *A. muciniphila* and *Faecalibaculum rodentium* species (**Fig. 6E**). Within both microbiome contexts, bile acids were associated with the 1-wk timepoint, suggesting depletion over time through the course of EAE and/or with extended time on experimental diets, although the ASVs associated with each time point differed dramatically between the two microbiomes (**Fig. 6D** and **F**). Taken together, these data identify microbial and metabolic signatures of diet and disease state, which differ between gut microbiome contexts.

**Figure 6.**
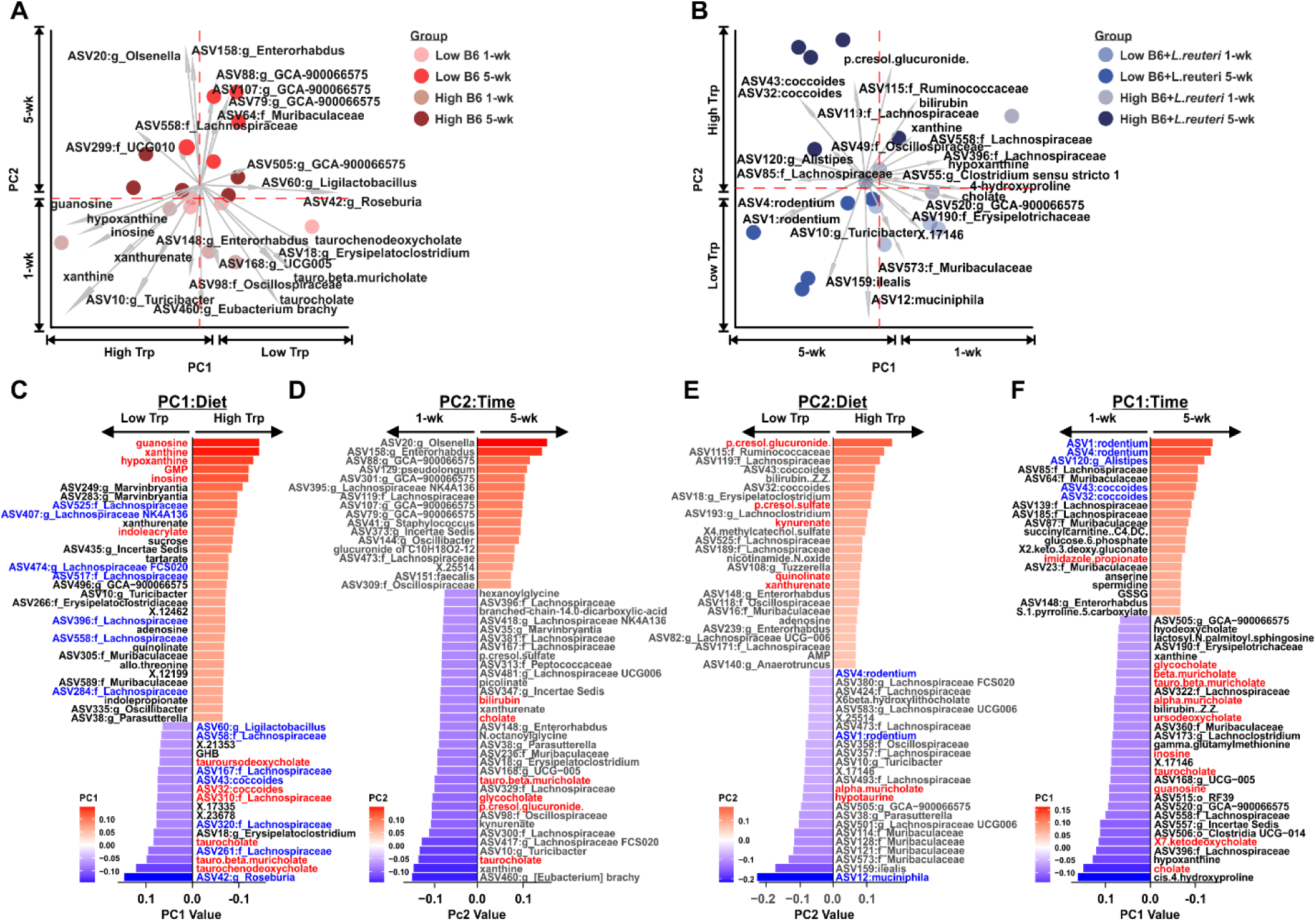
Bile acids, tryptophan, and associated microbiota delineate effects of diet and time in a microbiome-specific manner. Microbiome and metabolomic datasets were subset by gut microbiome configuration (B6 or B6 + *L. reuteri*) and integrated using jRPCA. Emperor biplot of integrated sample clustering based on multiomic variation, in the B6 (**A**) or B6 + *L. reuteri* (**B**) gut microbiomes, where vectors indicate the contribution of top 25? microbial and metabolic features to principal components. Top 50 features that delineate samples along principal component (PC) 1 and 2 in the B6 (**C** and **D**) or B6 + *L. reuteri* (**E** and **F**) gut microbiomes, segregating samples by dietary context or time.

### Identification microbial and metabolic signatures predicting EAE severity

To identify top microbial and metabolic biomarkers of EAE severity in our total dataset, we used Random Forests regression to predict EAE cumulative disease score (CDS, i.e. the sum of all daily scores) as a continuous variable from 1) the FL16S microbiomics dataset, 2) the metabolomics dataset, or 3) integrated microbiomic and metabolomic data. Features contributing to model performance were evaluated by their contribution to node purity and percent increase mean square error (MSE). Features of the microbiota-only model most highly contributing to performance included *Anaerotruncus (Ruminococcaceae),* and several *Lachnospiraceae,* and *Muribaculaceae* ASVs, as well as minor but significant contributions of *A. muciniphila*, *L. reuteri* (ASV74), *L. murinus*, *L. johnsonii*, *F. rodentium*, *Roseburia,* and *B. coccoides* (**Fig. 7A**). However, overall model performance was modest, with only a 0.599 correlation (p<0.01) between predicted and actual CDS, accounting for only 29.7% total variance with a 17.17 root MSE (**Fig. 7B**). For models trained on the metabolomics-only dataset, the top 30 features most contributing to overall performance mostly included tryptophan-associated metabolites, as well as those involved in NAD+ metabolism, and bile acids (**Fig. 7C** and **D**). Notably, the predictive capacity was higher (0.834, p<0.001) when using metabolomics features compared with ASV features, with 59.9% variance accounted for in the model and a predicted root MSE of 12.9 (**Fig. 7E**). Interestingly, in models built on joint metabolomics/ASV data, the top 30 features contributing to predictive capacity were dominated by metabolites, which largely overlapped with the metabolite-only trained regression models, and included not only microbially-produced indoles like 3-indoleglyoxylic acid, but also p-cresols (**Fig. 7F** and **G**).

**Figure 7.**
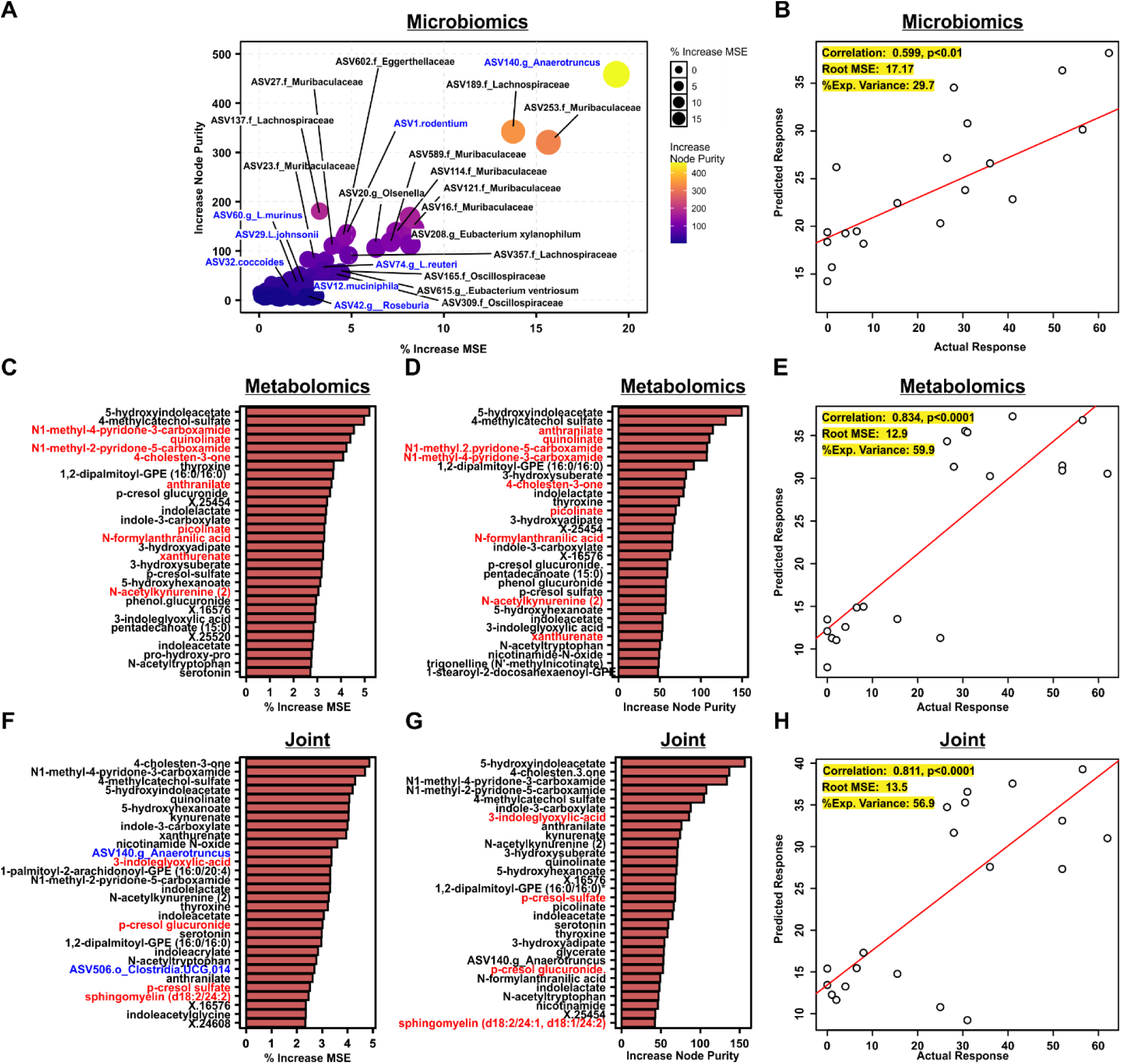
Systemic metabolites are better predictors of disease severity than the composition of the gut microbiome. (**A**) X-Y scatter plot reflecting total ASV variables of importance by % increase in mean squared error (MSE) and increase in node purity as determined by a Random Forest regression model trained on total ASV level FL16S abundance data to predict cumulative disease score (CDS), with a scatter plot of actual versus predicated values in (**B**). Top 30 metabolites as variables of importance by % increase in MSE (**C**) and increase in node purity (**D**) determined by a Random Forest regression model trained on total metabolomics data to predict CDS, with a scatter plot of actual versus predicated values shown in (**E**). Top 30 metabolites as variables of importance ranked by % increase in MSE (**F**) or increase in node purity (**G**), determined by a Random Forest regression model trained on joint FL16S and metabolomics data, with a scatter plot of actual versus predicated values shown in (**H**). Model performance was assessed by correlation between actual and predicted values, root mean square error, and the % explained variance. All models were trained using 1000 trees and optimized for feature selection per split and minimum samples per node.

Sphingomyelin metabolites were also identified as disease-predictive features (**Fig. 7F** and **G**), likely reflecting biomarkers of ongoing myelin damage in the CNS. Interestingly, two ASVs were included in the top 30 of the joint model: ASV140 (*Anaerotruncus)*, and ASV506 (Clostridia UCG.014) (**Fig. 7F** and **G**). Performance of the joint model was also comparable to the metabolite-only trained regression model (**Fig. 7H**). These data suggest that in this experimental model of dietary tryptophan modulation, circulating metabolites are a better predictor of disease severity than is the composition of the gut microbiome.

### Functional validation of metabolic signatures predicting EAE severity

Given our identification of p-cresols and indoles as associated with the B6 + *L. reuteri* microbiome (**Fig. 6**) and their value in predicting disease severity (**Fig. 7D-H**), we next sought to determine if these metabolites were sufficient to modulate CNS autoimmunity. We selected p-cresol, given the strong association with MS [48, 54] and our own studies showing that *L. reuteri* directly produces p-cresols and that these metabolites are tryptophan-dependent [59]. Notably, we selected unmodified p-cresol as our treatment, since this metabolite is normally produced by gut bacteria and subsequently sulfated or glucuronidated by the host to generate p-cresol-sulfate or p-cresol-glucuronide [64], both of which we detected in our metabolomic analyses. We also selected the novel, *L. reuteri*-produced and tryptophan-dependent metabolite, 3-indoleglyoxylic acid, a.k.a. indole-3-glyoxylic acid (IGoxA), based on robust clustering within the tryptophan network jointly with *L. reuteri* and p-cresols (**Fig. 5B** and **E**), and identification as a top biomarker of elevated disease severity (**Fig. 7F** and **G**).

**Figure S1.**
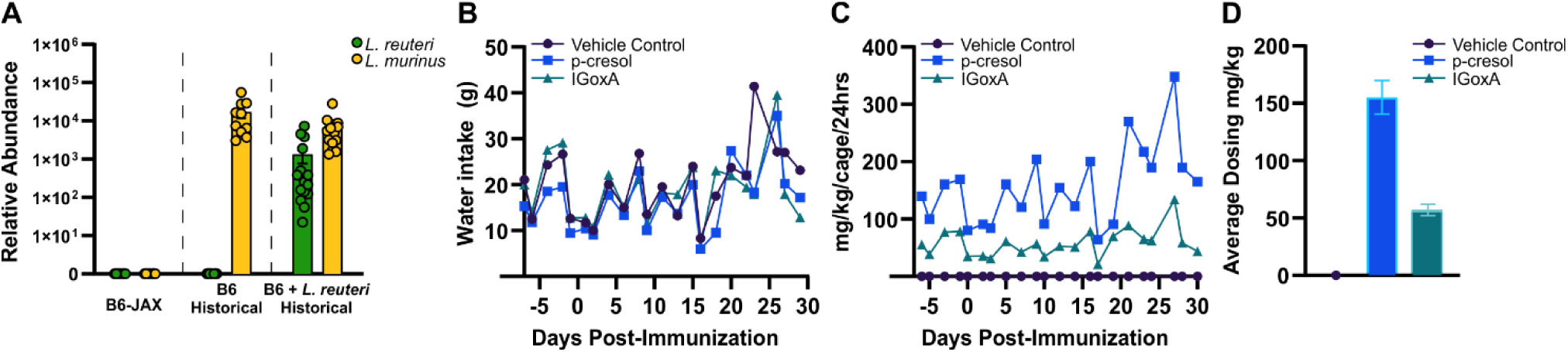
Identification of a colony of C57BL/6J mice that are naturally devoid of *L. reuteri* and *L. murinus*. (**A**) Relative abundance of *L. reuteri* and *L. murinus* in C57BL/6J mice from room AX29 distributed by the Jackson Laboratory was determined by species-specific qPCR and normalized by eubacterial pan-specific qPCR abundance. Historical B6 and B6 + *L. reuteri* abundance data was included as species-specific positive controls. (**B**) Water intake per cage of 5 mice. (**C**) Cage dosing in mg of metabolite per kg body weight. (**D**) Average experimental dosing per mouse in mg of metabolite per kg body weight. Water bottles and mouse weights were taken every other day.

To establish the sufficiency of IGoxA and p-cresol to modulate CNS autoimmunity, we provided each metabolite to mice via drinking water, followed by assessment of EAE severity. To minimize the effects of endogenous gut bacterial tryptophan metabolized, we leveraged a colony of C57BL/6J mice distributed by the Jackson Laboratory (B6-Jax), which we discovered to be naturally devoid of the *Lactobacillaceae* members, *L. reuteri* and *L. murinus* (**Fig. S1A**), which are major tryptophan metabolizers in the murine gut [59]. 7 week old B6-Jax mice were randomized to drinking water containing vehicle alone (filtered water), 5mM p-cresol, or 1mM IGoxA, with dose selected based on previous studies [46, 65–67]. Mice were treated for 1-week, followed by EAE induction, and continued treatment throughout the full 30-day disease course (**Fig. 8A**). Experimental water was refreshed weekly, and water intake and mouse body weights were measured every other day (**Fig. S1B-D**). Both p-cresol and IGoxA exacerbated EAE disease severity as compared to vehicle alone, as assessed by “classic” neurological symptoms of EAE (i.e. ascending paralysis; see Materials and Methods) (**Fig. 8B**). Interestingly, mice receiving the p-cresol treatment also displayed elevated symptoms of axial rotary EAE (AR)-EAE, an atypical EAE presentation consisting of ataxia and rotational movements (see Materials and Methods), primarily at disease onset (**Fig. 8C**). Cumulative disease score, the sum of all classic EAE scores over a 30-day disease course, was also elevated by both p-cresol and IGoxA treatment (**Fig. 8D**). Weight loss was not seen with treatment in the initial 1-week period prior to EAE induction, suggesting a lack of toxicity of the metabolite treatments; however, increased weight loss, as an indicator of disease severity, was observed after EAE onset with both metabolite treatments compared to the control group (**Fig. 8E**).

**Figure 8.**
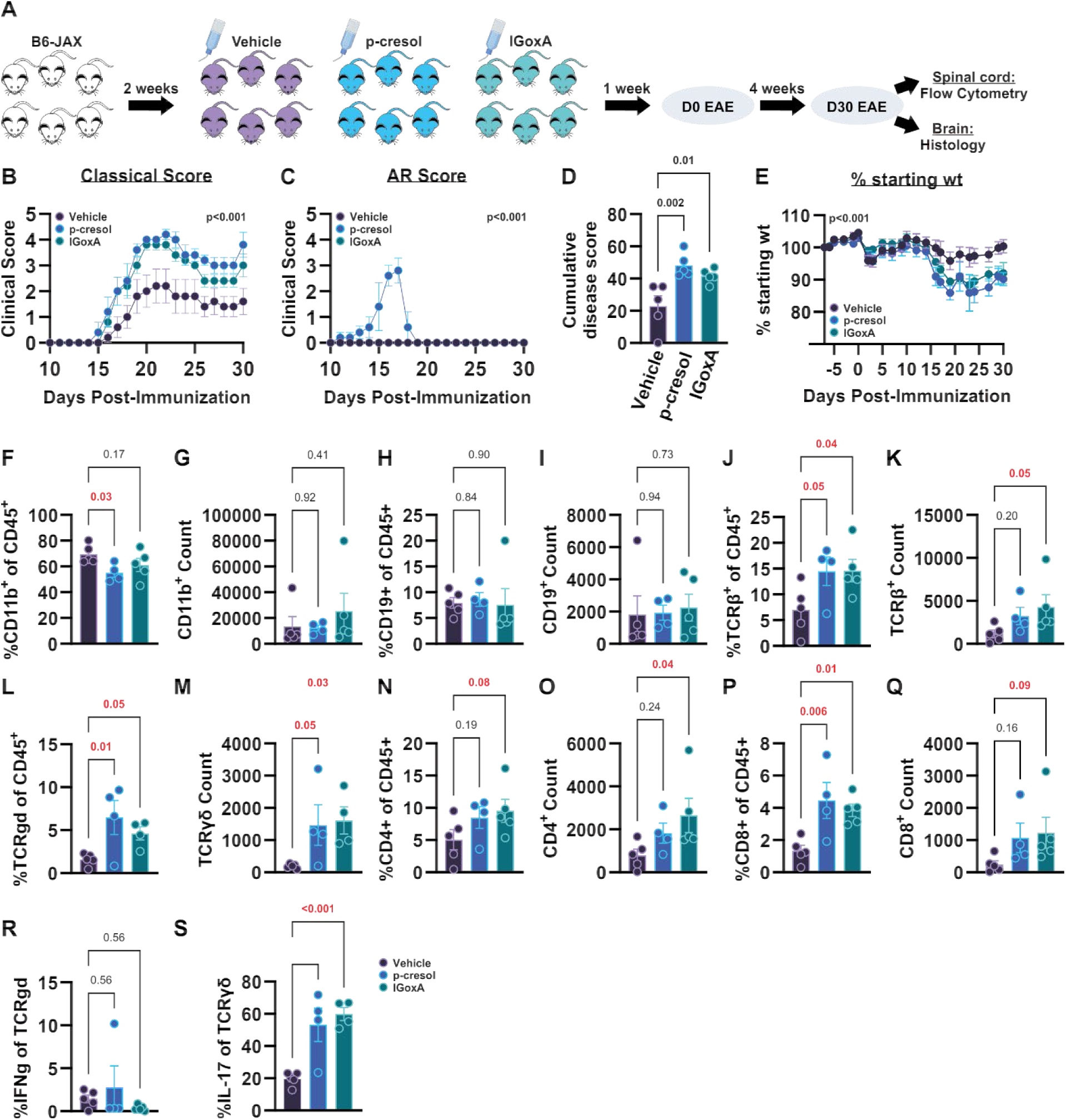
Indole and cresol metabolites are sufficient to exacerbate CNS disease pathogenesis, promoting proinflammatory T cell responses. (**A**) Schematic of experimental model depicting timeline for metabolite treatment with 5 mM p-cresol, 1 mM 3-indoleglyoxylic acid, (IGoxA), or vehicle control in the drinking water, relative to EAE induction in B6 Jax mice at 8 weeks of age (n=5M/group). EAE disease courses for each metabolite treatment are represented as mean daily clinical score with overall significance determined by Friedman’s parametric two-way ANOVA for (**B**) classical EAE scoring and (**C**) AR-EAE scoring, assessing the time*treatment interaction effect. CDS representing the total sum of all daily EAE scores over a 30 day disease course for metabolite treatment groups and is shown in (**D**), with significance evaluated using Mann-Whitney non-parametric test. (**E**) % starting weight represents mean daily percentage within each group with overall significance determined by Friedman’s parametric two-way ANOVA, assessing the time*treatment interaction effect. At day 30 post EAE induction, spinal cord-infiltrating leukocytes were isolated and analyzed by flow cytometry. Frequencies of the major leukocyte populations within the CD45^+^ population and total cell counts of (**F** and **G**) CD11b^+^, (**H** and **I**) CD19^+^, (**J** and **K**) TCRβ^+^, (**L** and **M**) TCRγδ^+^, (**N** and **O**) CD4^+^ T cells, and (**P** and **Q**) CD8^+^ T cells. Cytokine production as a frequency of TCRγδ^+^ T cells is shown for IFNγ (**R**) and IL-17 (**S**). Significance was determined by one-way ANOVA at p_adj_ ≤ 0.05.

To assess the CNS-relevant immunological impact of p-cresol and IGoxA treatment, immune cells infiltrating into the spinal cord of diseased mice were analyzed via flow cytometry. There were no major changes in CD11b^+^ microglia/myeloid cells or CD19^+^ B cells between treatment and control, aside from a slight decrease in the frequency of CD11b^+^ cells in p-cresol-treated mice (**Fig. 8F-I**). However, there was a greater frequency and number of both TCRβ^+^ and TCRγδ^+^ T cells driven by both metabolites (**Fig. 8J-M**). While IGoxA led to an increased frequency and number of both CD4^+^ and CD8^+^ T cells, p-cresol treatment resulted in only an increase in the frequency of CD8^+^ T cells (**Fig. 8N-Q**). Importantly, while not impacting IFNγ production, both metabolites led to a 2-3 fold expansion in the frequency of IL-17-producing γδ T cells (**Fig. 8S**). Taken together, these data establish that the bacterial metabolites p-cresol and IGoxA not only serve as biomarkers of EAE severity, but are sufficient to exacerbate CNS autoimmunity.

We and others have previously shown that atypical AR-EAE symptomatology is associated with inflammation and demyelination in the cerebellum rather than the spinal cord [68, 69]. To assess this possibility, we performed semi-quantitative histologic assessment of inflammation and demyelination in the cerebellum using H&E and Luxol Fast Blue (LFB) staining, respectively [68]. We found that p-cresol treatment elevated inflammation in the cerebellum (**Fig. 9A** and **B**), as expected from AR-EAE presentation (**Fig. 8C**), although demyelination was not significantly different (**Fig. 9C** and **D**). Surprisingly, we found that IGoxA treatment also elevated cerebellar pathology (**Fig. 9A-D**), in the absence of AR-EAE clinical signs (**Fig. 8C**). Taken together, these data suggest that, in addition to exacerbating clinical severity of EAE and CNS inflammation, both p-cresol and IGoxA may promote cerebellar pathology, which is prominently seen in pwMS [70].

**Figure 9.**
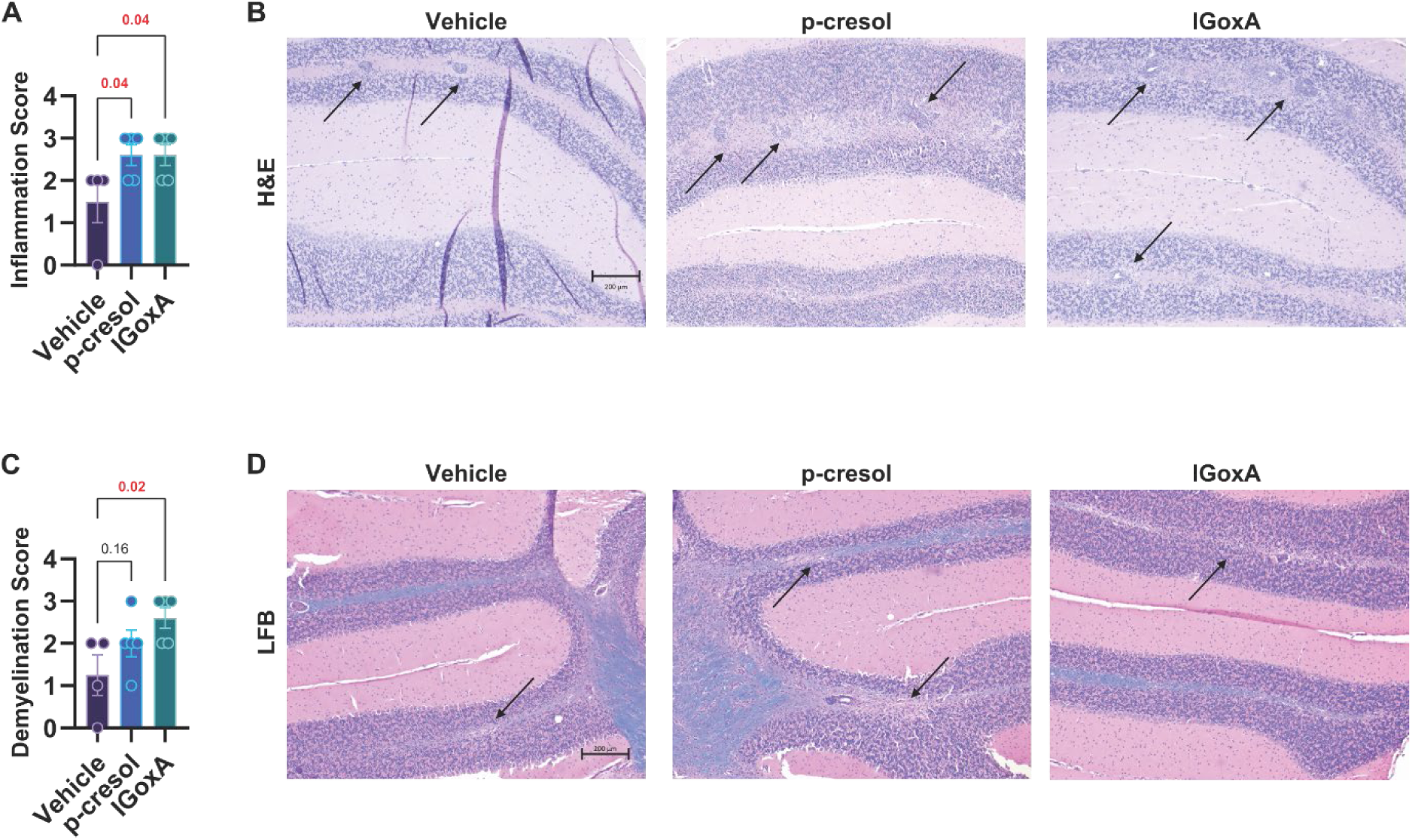
Indole and cresol metabolites promote cerebellar brain pathology. EAE and metabolite treatments were conducted as outlined in Figure 8A. At day 30 post EAE induction, brains were collected and processed for staining with H&E with or without LFB. Histopathologic evaluation of control, p-cresol, and IGoxA (n=4-5M/group) was performed as described in Methods. (**A** and **B**) Cerebellum inflammation scores by strain and corresponding H&E representative images. (**C** and **D**) Cerebellum demyelination scores by strain and corresponding LFB with H&E representative images. Cerebellum images (**B** and **D**) were captured at 5× objective. Scale bar: 200 μm. For all images, the arrows mark regions of inflammatory infiltrates or demyelination. Significance of differences was determined by ordinary 1-way ANOVA, with Fishers LSD multiple-comparison test at p ≤ 0.05.

## DISCUSSION

The gut microbiota is a putative environmental risk factor for modulating the development, progression, and/or severity of MS [10–12]. One key mechanism whereby the gut microbiota are postulated to influence autoimmune disease is the through the production of immunomodulatory metabolites, functioning locally in the gut, or at distal sites, including the CNS [71]. Dietary input is also thought to impact CNS autoimmunity and reshape the composition of the gut microbiome, altering the profile of bacterial and host derived systemic metabolites [72, 73]. The specific contribution of, and interplay between, the gut microbiota, diet, and production of immunomodulatory metabolites in MS is less clear. Here, we leveraged a longitudinal study design, in the EAE model of MS, using defined baseline gut microbiomes with layered dietary intervention, coupled with profiling of the gut microbiome and systemic metabolites to begin to define these interactions as relevant to disease. Using this approach, we found that a gut microbiome colonized with *L. reuteri* is more dynamic, exhibiting more significant changes over time, through the course of disease progression, and with longer-term dietary intervention, particularly in the context of high tryptophan, when compared to a microbiome lacking this species. Integration of metabolomic and microbiomic data, revelated unique clusters of metabolites and microbiota, that were enriched in distinct metabolic pathways, and sufficient to segregate samples by diet and timepoint within each microbiome context. Moreover, Random Forest modeling revealed that the abundance of systemic metabolites, functioned as more robust predictors of disease severity than did the gut microbiota, and identified cresol and indole tryptophan-associated metabolites among the top features contributing to overall model performance. Direct treatment with IGoxA and p-cresol in the EAE model, led to exacerbation of disease, eliciting increased inflammation and demyelination in the CNS. These data highlight the utility of leveraging a multi-omics approach to identify drivers of CNS autoimmunity and identify the baseline composition of the gut microbiota as a significant modifier of response to dietary intervention strategies.

Multiple case control studies have shown that pwMS have a distinct gut microbiome as compared to healthy controls [16–18]. Determining whether these changes predate disease onset, and therefore represent a risk factor for developing disease, or occur through the course of disease progression, representing either signatures of worsening disease severity or disease symptomology, is difficult to ascertain within human population without costly and lengthy longitudinal or prospective studies.

Consequently, the EAE model of MS represents an attractive alternative, within which the baseline gut microbiome can be tightly controlled and experimentally manipulated. Here, using a longitudinal study design, we found that with short-term dietary intervention, the baseline gut microbiome was a more significant driver of altered gut microbial composition than was diet alone and that these changes could persist through the course of disease. Interestingly, a microbiome lacking *L. reuteri* was relatively stable over time (**Fig. 4A-D**), exhibiting minimal changes directly following dietary intervention (**Fig. 2A, B**, and **F**) which persisted to the 5-week post-diet timepoint (**Fig. 3A, B**, and **F**). By contrast, independent of diet, *L. reuteri* colonization led to increased *A. muciniphila* abundance persisting through the course of EAE (**Fig. 2E** and **3E**). Interestingly, in *L. reuteri-*colonized mice fed a high tryptophan diet, a diet × microbiome configuration we have previously shown to be associated with elevated disease [59], *A. muciniphila* abundance was lower (**Fig. 3G**). These data are reminiscent of *A. muciniphila* abundance profiles in pwMS where this species is globally reported to be increased as compared to healthy controls, but diminished as associated with progression of disease [16–18]. One possible explanation for this paradoxical relationship between *A. muciniphila* abundance and CNS autoimmunity, is that this species is broadly beneficial and therefore host-selected in pwMS and similarly in mice harboring *L. reuteri*, but through the course of disease progression with worsening severity, abundance drops. These data suggest that the altered *A. muciniphila* abundance observed in pwMS may in part be dependent on the baseline composition of the gut microbiota, consistent with the inter-individual variation in *A. muciniphila* abundance observed in those studies.

A marked depletion of SCFA-producing *Lachnospiraceae NK4A136* was observed both at the 1-week and 5-week post-dietary intervention timepoints associated with both a high tryptophan diet and *L. reuteri* colonization (**Fig. 2I** and **3I**). This finding is consistent with our previous study demonstrating that *L. reuteri* actively depletes butyrate-producing microbiota to exacerbate CNS disease pathogenesis [60]. Of note, depletion of *Lachnospiraceae NK4A136* was not observed in a tryptophan depleted diet, even in the presence of *L. reuteri*, suggesting that the interaction between *L. reuteri* and SCFA-producers may be diet dependent. These data are consistent with other reports suggesting that SCFA exposure and dietary fructose, lead to a competitive fitness advantage for *L. reuteri* in the gut, which also highlight a dietary dependent relationship between *L. reuteri* and the microbiota that produce SCFAs [74, 75]. More broadly, one of the most consistent hallmarks of the MS gut microbiome is a depletion of known SCFA-producing microbiota, including *Butyricimonas*, *Bacteroides*, *Lachnospiraceae*, and *Eubacterium* [36–38, 76, 77] and their resulting metabolites [35–38, 40–42, 77, 78]. Our data suggest that this effect may be influenced by the combined effect of the baseline gut microbiome and dietary factors, potentially predates disease onset, and is sustained through the course of disease. *Blautia* (a *Lachnospiraceae* member known for SCFA-production) as a genus or individual species are increased in pwMS [19, 26, 79, 80], associated with increased disease severity, and responsive to DMT usage [16]. In our present study, we show that while *Blautia coccoides* expands throughout the course of EAE as a function of high dietary tryptophan and *L. reuteri* colonization. These data are also highly concordant with the profile seen directly in pwMS, where *Blautia* is associated with elevated severity of disease.

Several studies that have shown differences in *Lactobacillaceae* directly in pwMS [16, 17, 19, 81]. While lower in relapsing remitting MS (RRMS) compared with healthy controls, *Lactobacillaceae* are positively associated with disability [19] and are increased in abundance compared to individuals with progressive MS (PMS) [16]. Interestingly, *Lactobacillaceae* are also associated with worsening disease in PMS [17] and the *Lactobacillus* genus is positively associated with MS risk by Mendelian randomization [81]. We and others have previously shown that stable colonization by the gut commensal bacterium *L. reuteri* is sufficient to exacerbate EAE [56–58]. In contrast, other studies have shown that continuous treatment with a high dose of *L. reuteri* has the opposite effect [82, 83]. This suggests that the effect of *L. reuteri* differs between natural colonization vs. probiotic-like treatment. Another possibility is that the different strains of *L. reuteri* used in these studies could exert different effects, which is strongly supported by a recent study from the Walter group demonstrating strain-dependent effects of *L. reuteri* on EAE which correlated with the strains’ capacity to activate the AhR [57]. Given these potentially conflicting effects, in this study, we focused our integrative omics analysis on identifying the strongest biomarkers of CNS disease severity agnostic to the abundance of *L. reuteri* itself, which turned out to be metabolites rather than microbes, two of which we functionally validated in the EAE model. Our findings suggest that the collective output of the microbiota may be more important than individual members in terms of driving disease outcome. Interestingly, the abundance of *L. reuteri* itself did not serve as a robust biomarker of disease severity, while an ASV belonging to the *Anaerotruncus* genus was the best microbial predictor. These findings further reinforce our previous findings that *L. reuteri* may exert its effects on the host indirectly via other members of the gut microbiota, which in this study we show it affects to a much greater extent in the presence of dietary tryptophan. This raises the intriguing possibility that indoles and p-cresols may be produced by gut microbiota as antimicrobials that provide a competitive advantage in the gut [84].

In our study, we integrated microbiomic and serum metabolomics datasets, revealing clusters of metabolites and associated microbiota enriched in distinct metabolic pathways. Network analysis of the bile acid cluster identified members of the *Clostridia* class, including *Lachnospiraceae*, *Lachnospiraceae NK4A136*, and *Oscillospiraceae*, as well as *Olsenella* and *Muribaculaceae* members, as closely associated. Notably, prominent secondary bile acids known to be produced by these microbiota were also present in this cluster (7-ketodeoxycholate, 3-dehydrocholate, deoxycholate, hyodeoxycholate, and ursodeoxycholate) in addition to the host primary and conjugated bile acid from which they are derived (cholate, glycolate, glyco-beta-muricholate,, tauroursodeoxycholate, glycochenodeoxycholate, beta-muricholate, alpha-muricholate, chenodeoxycholate, and glycoursodeoxycholate) (**Fig. 5B, C, D** and **F**) [85]. Moreover, bile acids were among the top 50 features of importance segregating both B6 and B6 + *L. reuteri* by diet and/or timepoint, where elevated bile acids were more robustly associated with the B6 microbiome, a low tryptophan diet, and diminished through the course of disease (**Fig. 6**), suggesting a protective role for bile acids and/or microbiota that produce them. Furthermore, bile acids were among the top metabolomic features, particularly 4-cholesten-3-one, contributing to predictive capacity of disease severity within Random Forest models (**Fig. 7**). Bile acids are thought to broadly exert an anti-inflammatory effect. Both primary and secondary bile acids are reduced in circulation in pwMS, and supplementation safely restores plasma levels [33, 86–88]. However, the precise metabolites and microbiota relevant to disease are less clear, and matched microbiomic/metabolomic datasets from pwMS and/or through the course of supplementation may be warranted to define relevant intervention strategies.

Tryptophan-associated metabolites are also altered in the serum, plasma, CSF, and urine of pwMS. Tryptophan itself is depleted in the serum or plasma of pwMS [46–48]. Levels of 5-hydroxytryptophan, kynurenine, kynurenate, quinolinate, xanthurenate, and the bacterial metabolites, indole-3-lactate (ILA) and indole-3-propionate are also depleted in circulation and increases in indole-3-acetate (IAA) and 3-hydroxyanthranilate have been found [46, 48, 49, 51, 89]. More recently, elevated levels of the bacterial oxidative metabolites of tryptophan, in particular IAA, which is elevated in MS, have been implicated in demyelination, whereas provision of the reductive metabolite ILA, which is depleted in pwMS, function in an anti-inflammatory fashion to promote remyelination, with promising preclinical results [90]. This suggests that among the bacterial tryptophan-metabolites collectively referred to as “indoles”, individual metabolites can function in unique and sometimes opposing ways. Given the increased focus on direct indoles as therapeutic intervention strategy in pwMS, more precisely defining the microbiota involved and their downstream metabolic products likely to be resulting from such supplementation strategies is warranted, particularly given the often-contradictory impact of various bacterial produced indoles on CNS autoimmunity. In our study, network analysis of the cluster enriched in metabolites contributing to the tryptophan pathway suggested more taxonomic diversity represented among closely associated microbiota than was observed within the bile acid cluster (**Fig. 5B, C, E**, and **F**). Tryptophan associated metabolites were also among the top 50 features contributing to separation of B6 + *L. reuteri* samples by diet (increased with high tryptophan) (**Fig. 6**) and among the top 30 features contributing to Random Forest predictive modeling of disease severity (**Fig. 7**). Interestingly, IGoxA contributed highly to both indicators of model importance (node purity and % increase in MSE), suggesting this indolic metabolite is a strong predictor of clinical disease severity. We have previously shown that IGoxA is directly produced by *L. reuteri* in monoculture, robustly activates the aryl hydrocarbon receptor, enhances T cell production of IL-17, and is produced through the aromatic amino acid aminotransferase (ArAT) tryptophan degrading pathways, which also produces IAA and ILA [59]. Direct treatment with IGoxA was sufficient to exacerbate disease within the EAE model, eliciting enhanced inflammatory T cell production and demyelination. We observed not only increased frequency of CD4^+^ and CD8^+^ T cells but also an increase in γδ T cells and their IL-17 production. These data are consistent with our previous findings demonstrating an *L. reuteri* and tryptophan dependent increase in γδ T IL-17 production [59]; however, in this case this effect was directly mediated by individual tryptophan-dependent and *L. reuteri* produced metabolites. While we have not demonstrated a direct functional role for IL-17 producing γδ T cells, these findings suggest that at the very least this T cell subtype, which is already known to promote EAE [91], is sensitive to microbial metabolites, consistent with findings in other studies [92]. Moreover, While IGoxA has not been previously shown to be altered directly in pwMS, these data support the notion that diverse metabolites from the bacterial ArAT pathways that can either exacerbate disease, as found here for IGoxA and previously reported for IAA [93], or support remyelination, as has been shown for ILA treatment [90]. Importantly, future studies can determine whether these indole-dependent effects on specific host cell types are driven by differential activation of the major known receptor for these xenobiotic molecules, the aryl hydrocarbon receptor (AhR), or other mechanisms.

Cresol metabolites were also among the top 50 features segregating the B6 + *L. reuteri* microbiome samples by dietary context (**Fig. 6**) and among the top features contributing to overall predictive modeling of clinical severity (**Fig. 7**). This was an unexpected finding given that cresols are canonically associated with bacterial phenylalanine and tyrosine metabolism [94]. However, there is evidence to suggest that bacteria can also utilize tryptophan to produce cresol which is supported by our integrated analysis here, clustering cresols with tryptophan metabolites and the microbiota that produce them (**Fig. 5**) [95]. Importantly, cresols are increased in pwMS and other neurologic disorders and are considered as potentially neurotoxic [48, 54, 94, 96]. We have previously reported that both p-cresol sulfate and p-cresol glucuronide are elevated in a tryptophan dependent manner, consistent with multi-omics analysis here, and that cresols are produced directly by *L. reuteri* in monoculture in a tryptophan-dependent fashion [59]. In this study, we demonstrate that treatment with p-cresol exacerbates classic neurological symptoms of disease in the EAE model, and also elicits a transient but severe atypical AR-EAE presentation primarily with ataxia early in the disease course as well as enhancing inflammatory T cell production in the spinal cord and inflammation in the cerebellum. Interestingly, the immunological phenotype was similar to that observed with IGoxA treatment, with a noted expansion of CD8^+^ T cells, TCRγδ T cells, and TCRγδ IL-17 production in the spinal cord of disease mice by day-30 post-induction. Given that ataxia resolved by day-20, additional studies will be needed to determine the role of cresols in reshaping the immune response and the CNS environment, particularly early in disease. Cresols have been shown to induce axonal damage, and to impair oligodendrocyte differentiation reducing myelin production *in vitro* [54, 97], which would be consistent with enhanced demyelination observed with p-cresol treatment *in vivo* in our study. Moreover, immune recognition of p-cresol sulfate has been reported to be similar to that of myelin basic protein, suggesting that cresols may function as myelin autoantigen mimics [98]. These data underscore the potential role of cresols in driving MS disease pathogenesis.

Taken together, our data highlight the utility of leveraging a multi-omics integrated approach coupled with a longitudinal study design to understand the microbiomic, metabolomic, and immunological drivers of MS, which we leveraged here to prioritize metabolites for direct preclinical evaluation. Moreover, our findings underscore the importance of the baseline gut microbiota in modulating response to dietary intervention strategies as influencing immunomodulatory host and bacterial derived metabolites.

## MATERIALS AND METHODS

### Lactobacillaceae isolation

*Lactobacillaceae* were isolated as previously described [59]. *Limosilactobacillus reuteri* (*L. reuteri*) was isolated from the cecal contents of a PWD/PhJ mouse, as follows. Total cecal contents were incubated overnight anaerobically at 37°C, with 200 rpm shaking resuspended in 50 ml DeMan, Rogosa and Sharpe (MRS; Thermofisher, Inc, USA) medium supplemented with 0.25g/L L-cysteine (Sigma, USA), 20µg/ml vancomycin (Sigma, USA) adjusted to pH 5. Cultures were plated onto agar medium of the same formulation and incubated anaerobically overnight at 37°C. Isolated colonies were cultured overnight in MRS medium, cryopreserved, and screened by qPCR with species-specific primers [56]. Positive isolates for each species were cultured in vancomycin-free medium followed by repeat screening by qPCR. For colonization studies, gavage stocks were prepared using three isolates per species grown to log-phase and adjusted to OD600=0.5 with fresh medium. Colony forming units (CFU) of gavage stocks were quantified by standard serial dilution and plating.

### Animals and microbiota transplantation

Gut microbiota transplantation was performed as previously described [56]. Briefly, cecal contents were cryopreserved from B6 or PWD mice under anaerobic conditions, in 20% glycerol in Hungate tubes and stored at -80°C until use. 4-5 week old C57BL/6J germ-free (GF) mice were purchased from the National Gnotobiotic Rodent Resource Center at University of North Carolina School of Medicine (Chapel Hill, NC, USA). Upon arrival, GF mice were immediately inoculated by gastric gavage with 100µl of cryopreserved PWD or B6 cecal content, under a laminar flow hood. For mice receiving the B6+*L. reuteri* microbiota, 100µl of the B6 microbiota supplemented with 100µl *L. reuteri* at 10^9^ CFU was used. Colonized ex-GF mice served as breeding pairs for a vertical gut microbiota transmission model.

Conventional SPF C57BL/6J mice were purchased from Jackson Laboratory, immediately unpacked under a laminar flow hood upon arrival, and rested for 1-week prior to experimentation. All animals were maintained under barrier conditions with sterilized food, water and caging, with minimal handling. The experimental procedures used in this study were approved by the Animal Care and Use Committee of the University of Vermont.

### Induction and evaluation of EAE

EAE was induced using the 2×MOG35-55/CFA protocol in either exGF-B6 GMT recipients or C57BL/6J (Jackson Laboratory) [99]. On day 0 (lower flank) and day 7 (upper flank) mice were injected subcutaneously with 50 µl per flank of 0.1 mg of myelin oligodendrocyte glycoprotein peptide 35-55 (MOG35-55) peptide (New England Peptide, Inc. MA, USA) emulsified in PBS and 50% complete Freund’s adjuvant (CFA; Sigma, USA) supplemented with an additional 4 mg/ml Mycobacterium tuberculosis H37Ra (Difco, USA). Beginning on day 10, mice were scored as follows for “classic” signs of EAE: 1 – loss of tail tone, 2 - loss of tail tone and weakened hind limbs, 3 - hind limb paralysis, 4 - hind limb paralysis and incontinence, 5 - quadriplegia or death. For axial rotatory (AR)-EAE, the mice were scored as follows: 0, asymptomatic; 1, slight head tilt; 2, pronounced head tilt; 3, inability to walk in a straight line; 4 mouse is moving/lying on its side, will continuously fall to its side after being made to stand; 5, mouse rolls or spins continuously. Differences in disease course and weights were evaluated using Friedman’s non-parametric two-way ANOVA [99], using the treatment by time interaction term to signify a significant treatment effect. Cumulative disease score (CDS) is the sum of all scores over the 30-day disease course with significant differences determined using one-way ANOVA.

For dietary intervention studies, defined microbiome colonized mice were randomized to a low 0.02% or 0.8% high tryptophan diet (TD.200350 and TD.200352, Envigo Teklad Diets, Madison, WI) one week prior to EAE induction at 8-12 weeks of age. Diets were vacuum packaged, irradiated, stored at 4°C until use, and refreshed weekly in experimental cages. Weights were taken every other day for the duration of the experiment.

For metabolite treatments, C57BL/6J were randomized to control (sterilized water), 1 mM indole-3-glyoxylic acid (AK Scientific, Union City, CA), or 5mM p-cresol (Sigma, USA) one week prior to EAE induction at 8 weeks of age. Experimental water was refreshed weekly. Weights were taken every day.

### Flow cytometry

At day 30 post-EAE induction, mice were anesthetized with isoflurane and transcardially perfused with PBS. Spinal cords were removed, homogenized to generate a single cell suspension, and filtered with a 70 µm strainer followed by Percoll gradient (37%/70%) centrifugation and collection of the interphase. For intracellular cytokine staining, cells were stimulated with 20 ng/ml PMA, 1 µg/ml of ionomycin, and 1mg/ml brefeldin A (Golgi Plug reagent, BD Bioscience) for 4 hours. Cells were stained with the UV-Blue Live/Dead fixable stain (Thermofisher, USA) and antibodies against CD45, CD11b, CD19, TCRβ , CD4, CD8, and TCRγδ, (Biolegend, USA). For intracellular cytokine staining, cells were fixed, permeabilized with 0.05% saponin, and stained with anti-IL-17A and anti-IFNγ antibodies (Biolegend, USA). Cells were analyzed using an Cytek Aurora (Cytek Biosciences, USA). Spectral unmixing was performed with using single-color controls and autofluorescence correction from an unstained control. Data were analyzed using FlowJo software, version 10.8.1 (Tree Star Inc, Ashland, OR).

### CNS histopathology

At day 30 post-EAE induction mice were euthanized, and brain tissues was collected for histopathological assessment as previously described [99]. Briefly, the skull was removed and diffusion fixed in 10% neutral buffered formalin (Thermo Fisher Scientific). After fixation, the brain was extracted from calvaria and dissected into thirds (coronal orientation front brain, mid brain, and hind brain — including cerebellum and brain stem. Tissues were embedded in paraffin, sectioned coronally, and stained with LFB and/or H&E. Histopathological assessment was conducted in a blinded fashion. Regions were assessed for degree of inflammation and demyelination using a semiquantitative scale adapted from previous studies[99]. Inflammation was evaluated using H&E-stained tissues and scored as follows: 0, no inflammation; 1, few inflammatory cells scattered/small clusters; 2, organized clusters of inflammatory cells without significant extension beyond small lesions; 3, significant organized clusters of inflammatory cells with patchy infiltration of surrounding tissue, central involvement of larger lesions; and 4, extensive and dense infiltration of inflammatory cells affecting over half of the tissue. Extent of demyelination was evaluated using tissues stained with LFB + H&E and scored as follows: 0, no demyelination deep blue staining; 1, small, patchy area(s) of white matter pallor, no well-defined lesions; 2, defined area of white matter pallor forming isolated lesion(s); 3, confluent foci of white matter pallor with some spared areas; and 4, widespread white matter pallor affecting over ∼75% of the tissue.

### Full-length 16S (FL16S) sequencing

Fecal samples were collected as previously described [56, 59]. Individual mice were placed in sterile unbedded cages and allowed to defecate, fecal samples collected, and stored at -80°C. DNA was extracted using the QIAamp PowerFecal Pro DNA extraction kit (Qiagen, USA) and assessed via Nanodrop, Qubit, and agarose gel electrophoresis. Library construction and sequencing were performed at the Roy J. Carver Biotechnology Center, University of Illinois at Urbana-Champaign. Full length 16S (FL16S) amplicons were created using 2.5ng of genomic DNA according to the published PacBio 16S protocol. Briefly, PCR reactions were set up on ice in 25 µl volumes using barcoded primers with the following reagents: 12.5 µl Kappa Hotstart Ready Mix (KAPA Biosystems), 1.5 µl water, 3 µl (2.5 µM) 16S Forward primer (AGRGTTYGATYMTGGCTCAG), 3 µl (2.5 µM) 16S Reverse primer (RGYTACCTTGTTACGACTT), 5 µl DNA (0.5 ng/µl). PCR cycling parameters were: denature 95°C for 3 minutes, 25 cycles of 95°C for 30 seconds, 57°C for 30 seconds, 72°C for 60 seconds, and hold at 4°C. All products were measured with SpectraMax®Quant™ AccuClear™ Nano kit and run on an Agilent Fragment Analyzer for QC. Samples were pooled according to concentration and cleaned twice with 0.6 volumes of magnetic beads. The cleaned pool was checked on an Agilent Fragment Analyzer before library production. Amplicons were converted to a library with the SMRTBell Express Template Prep kit 2.0 (Pacific Biosciences, Menlo Park, CA). The library was sequenced on 1 SMRTCell 8M on a PacBio Sequel IIe using the CCS sequencing mode and a 15hs movie time. CCS analysis was done using SMRTLink V10.0 using the following parameters: ccs --min-passes 3 --min-rq 0.999, lima --ccs --different --split-bam-named. Demultiplexing of the PacBio library was done with SMRT link V11 (Pacific Biosciences, Menlo Park, CA).

### FL16S preprocessing and taxonomic assignment

Raw sequencing reads were analyzed using *DADA2* (version 1.34.0) in R (version 4.4.2) [100, 101]. Primer sequences and phiX reads were removed, reads filtered to retain a minimum length of 1000 and maximum length of 1600, denoised at a maximum number of errors (maxEE) of 2 and a quality score of 3. Reads were dereplicated followed by model-based error learning optimized for PacBio reads, denoising, and chimera removal using the consensus method. Taxonomy was assigned using the SILVA database (version 138.1) [102]. A phylogenetic tree was constructed by aligning unique sequences using *DECIPHER* (version 2.26.0) and generating a neighbor-joining tree with *phangorn* (version 2.12.1) using a maximum likelihood approach [103, 104]. The tree was optimized with nearest-neighbor interchange under a generalized time-reversible model and rooted using midpoint rooting [103–105]. Downstream analysis was conducted with *phyloseq* (version 1.50.0) [106]. Reads that did not align to the bacterial kingdom, *Cyanobacteria as* likely contaminants, and phyla falling below a mean prevalence of 1 (*Patescibacteria*) were removed.

### Diversity analysis

Alpha diversity (Shannon) was calculated using the plot_richness function in *phyloseq* (version 1.50.0) and statistical difference of the mean determined by Student’s t-test [107]. Unweighted Unifrac beta diversity was calculated using the distance function in *phyloseq* (version 1.50.0) and represented as a PCoA with statistical analysis of group differences using multivariate analysis of variance (adonis2) using *vegan* (version 2.6-8) [108, 109].

### Taxonomic bar graphs and differential abundance testing

Closely-related taxa were agglomerated using single-linkage clustering with the tip_glom() function in phyloseq. The AGNES algorithm was used with a numeric scalar of 0.1 as the distance threshold for merging taxa [110]. Taxa were filtered at a 2% prevalence threshold with a mean abundance threshold of 1 read per sample. Bar graphs are the top 11 most abundant taxa, excluding ASVs that do not assign to the rank of genus. Differential abundance of individual ASVs were conducted using LinDA (version 0.2.0), correcting for microbiome, diet, or both [61]. ASV level represents a taxonomic ‘best hit’ which was assigned using the format_to_besthit function in *microbiomeutilities* (version 1.00.17). Longitudinal analysis was conducted with the MicrobiomeStat (version 1.2.1) package which implements LinDA for feature level analyses [61].

### Metabolomics

Serum was collected from mice colonized with the B6 microbiome with or without *L. reuteri* fed a low (0.02%) or high (0.8%) tryptophan diet for 1 week or 5 weeks. Sera were stored at -80°C until shipment on dry ice to Metabolon Inc. (Durham, NC) for UPLC-MS/MS [111]. Proteins were precipitated with methanol with shaking for 2 min, followed by centrifugation and division into 5 fractions followed by removal of the solvent with a TurboVap, and overnight storage under nitrogen. Dried extracts were reconstituted for 1) acidic positive ionization of hydrophilic compounds: elution from a C18 column (Waters UPLC BEH C18-2.1×100 mm, 1.7 µm) with water and methanol, containing 0.05% perfluoropentanoic acid (PFPA) and 0.1% formic acid (FA), 2) acidic positive ionization of hydrophobic compounds: elution from the same C18 column using methanol, acetonitrile, water, 0.05% PFPA and 0.01% FA 3) basic negative ionization: elution from a separate C18 column using methanol and water with 6.5mM ammonium bicarbonate at pH8, and 4) negative ionization: elution from a HILIC column (Waters UPLC BEH Amide 2.1×150 mm, 1.7 µm) using a gradient consisting of water and acetonitrile with 10mM ammonium formate, pH 10.8. All methods used ultra-performance liquid chromatography (UPLC) and high resolution/accurate mass spectrometer with heated electrospray ionization and 35,000 mass resolution using an Orbitrap mass analyzer. Analysis used iterative mass spectrometry and data-dependent tandem mass spectrometry scans using dynamic exclusion and a range of 70-1000 m/z. Raw data was extracted for peak identification and quality assessment using Metabolon’s hardware and software with a proprietary library of 3300 authenticated purified standards.

### Integrated analysis

Serum metabolomics data was normalized by row median excluding zero values. FL16S data was subset to include only samples for which there was matched metabolomics data. Datasets were integrated using Joint Robust Aitchison PCA (Joint-RPCA) [63, 112] with a min_feature_frequency of 5 and max_interations of 15. For the total dataset, 26 samples were used in training and 10 for testing, split evenly between diet and microbiome groups. For analysis within each gut microbiome, 6 samples were using in training and 12 as a test set based on overall model performance. Resulting biplots and correlation matrices were imported into R (version 4.4.2) using *qiime2R* (version 0.99.6) for further analysis.

### Pathway enrichment analysis

The jRPCA correlation matrix was filtered to retain only ASVs as rows and metabolites as columns and clustered using *pheatmap* (version 1.0.1.2) using Euclidean distance and complete linkage. Clusters were automatically detected from row and column dendrograms by cutting the trees at fixed heights (25 for rows and 17 for columns) resulting in 7 distinct metabolite clusters and 11 ASV clusters. Metabolites were extracted from each cluster for pathway enrichment analysis with MetaboAnalyst (version 6.0) [113]. Analyses were performed using unique HMDB identifiers, with hypergeometric test as the enrichment method and relative-betweenness centrality as the topology measure, referencing the mouse Small Molecule Pathway Database (SMPD).

### Network analysis

Correlated ASVs and metabolites within the tryptophan or bile acid cluster were used to construct an undirected network using *igraph* (version 2.1.3) filtering data at a threshold of 0.99 where nodes represent metabolites or ASVs with color assignment based on taxonomic family. The network layout used a force-directed algorithm (layout_with_fr) and final visualization was performed using *ggraph* (version 2.2.1) where edge widths represent the strength of correlation and colors indicate the direction of correlation.

### Biplot variable importance

Biplots generated using jRPCA were imported into R (version 4.4.2) using *qiime2R* (version 0.99.6). The top 50 variables with the highest absolute loading values along component one or two (segregating samples by time or diet) were extracted and visualized as horizontal bar plots within each microbiome.

### Random forest modeling

Random forest regression models were constructed to predict cumulative disease score (CDS) from centered log ratio (CLR) transformed FL16S data, median scaled and log-transformed metabolomics data or the joint data using the *randomForest* (version 4.7-1.2) and *caret* (version 7.0-1) packages in R (version 4.4.2). Models were optimized using repeated 10-fold cross-validation (3 repeats) for the number of predictors sampled at each split (mtry) via random search (tuneLength =15) and nodesize for the lowest root mean square error (RMSE). Final models used the following parameters: FL16S (mtry =48, nodesize =1), metabolomics (mtry =62, nodesize =3), and joint (mtry =74, nodesize =1) with 1000 trees. The top 50 features contributing to RMSE and % increase in mean squared error (%IncMSE) were visualized as horizontal bar plots for each model.

## DECLARATIONS

### Ethics approval and Consent to participate

The experimental procedures used in this study were approved by the Animal Care and Use Committee of the University of Vermont under protocol ID PROTO202000037.

### Consent for publication

Not applicable

### Availability of data and materials

Microbiomic datasets generated and/or analyzed during the current study are available at the NCBI Sequence Read Archive repository (https://www.ncbi.nlm.nih.gov/bioproject/PRJNA1287598/). Metabolomic datasets generated and/or analyzed during the current study are available as relative abundance in supplementary table S29 and peak area in supplementary table S30. Raw spectral data are proprietary to Metabolon, Inc, and therefore are not available for deposit to a public repository.

### Authors’ contributions

T.L.M. and D.N.K. designed research; T.L.M., E.A.N, ,L.A.D., E.R.H., M.F.J.L performed research; T.L.M., C.M., D.M., G.R., R.K. and D.N.K. analyzed data; T.L.M. and D.N.K. wrote the paper; All authors reviewed the manuscript

## Acknowledgements

Dr. Gary Mawe, Dr. Jessica Crothers, Dr. Jon Boyson, and Dr. Cory Teuscher (University of Vermont) are acknowledged for helpful discussions and feedback.

